# Fibrotic-like collagen matrices as innovative 3D in vitro models for investigating the impact of pathological ECM on muscle regeneration in muscular dystrophies

**DOI:** 10.1101/2024.12.23.630059

**Authors:** Marie Camman, Naomi Nieswic, Pierre Joanne, Julie Brun, Alba Marcellan, Onnik Agbulut, Christophe Helary

## Abstract

Muscular dystrophies are characterized by impaired skeletal muscle contraction due to genetic mutations. Beyond the disruption of muscle function, the extracellular matrix (ECM) surrounding muscle fibers becomes fibrotic, further impeding tissue repair and function. In this study, we designed both healthy and fibrotic matrices to examine the impact of persistent fibrosis on muscle cell behavior. Fibrotic matrices were synthesized by 3D printing of dense collagen solutions in air, followed by slow gelation to yield non-porous, isotropic hydrogels. These matrices were subsequently crosslinked using EDC/NHS chemistry, resulting in a Young’s modulus of approximately 50 kPa. The behavior of C2C12 myoblasts cultured within the fibrotic matrices was compared to cells grown in healthy matrices. Our results showed that the fibrotic matrix had a detrimental effect on cell behavior. Myoblasts were unable to differentiate into mature myotubes, exhibited poor alignment, and suffered from hypoxia. Furthermore, these cells failed to proliferate, secreted inflammatory cytokines, and were unable to remodel their ECM. Using matrices possessing a single characteristic of the fibrotic matrix (i.e stiff or non porous), the predominant factor for each feature of the cell phenotype was determined. These findings underscore the detrimental effects of a fibrotic persistent ECM on muscle homeostasis. The development of these two distinct 3D muscle models, one representing healthy muscle and the other fibrotic, offers a valuable tool for investigating the pathophysiology of muscular dystrophy.

**Highlights:**

- Stiff and non-porous dense collagen matrices synthesized by 3D printing recapitulate the key characteristics of fibrosis observed in muscular dystrophies
- Fibrotic matrices negatively impact myoblast proliferation and differentiation into mature myocytes
- Fibrotic matrices impede the 3D organotypic organization of myocytes
- This 3D pathological model of skeletal muscle offers a valuable tool for investigating how persistent fibrosis influences the progression of muscular dystrophy

**Graphical Abstract:** 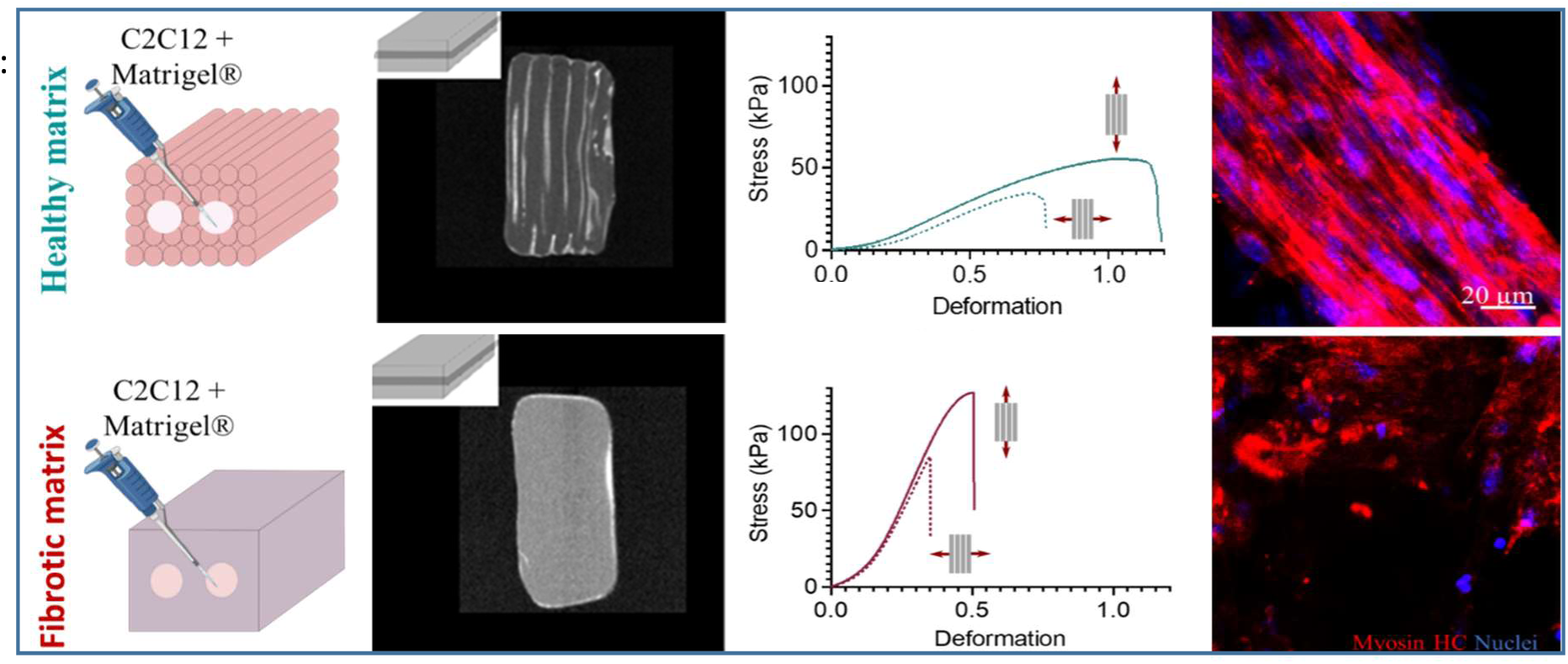

## 1. Introduction

Skeletal muscle is a key player in the body’s locomotion. The striated muscle is composed of fascicles of fibers responsible for muscle contraction, linked together by a specific extracellular matrix (ECM). The latter ensures the force transfer between cells and serves as mechanical support. It also directs cell growth, differentiation and muscle renewal [1]. The ECM surrounding fiber fascicles is mainly composed of type 1 and 3 collagen, is highly anisotropic and deformable with a Young’s modulus around 12-15 kPa [2].

The muscles of patients affected by muscular dystrophies are subjected to several adaptations and, among others, the extracellular matrix (ECM) surrounding muscular fibers undergoes a strong remodeling that participate to the impairment of muscle function [3]. This remodeling is mainly due to asynchronous cycles of degeneration and regeneration that provoke excessive accumulation of several ECM components. Type 1 Collagen is the major component of the accumulated ECM, which is involved in the pathological evolution toward fibrotic connective tissue formation [4]. The main features of such fibrotic connective tissue are high rigidity (Young’s modulus: 50-70 kPa), loss of anisotropy and reduced vascularization [5–7]. Moreover, the replacement of muscle cells by a fibrotic matrix will also alter muscle function such as contraction or regeneration properties [8]. For example, it has been demonstrated that persistent fibrotic matrix can affect mechanotransduction [9], myoblast activity [10] and fibro-adipogenic progenitors and myofibroblasts activation [11]. Because ECM is not an inert support for cells and has a bioactive role in the progression of muscular dystrophies, assessing its precise role seems to be of crucial importance. Especially because fibrosis may represent a very interesting therapeutic target [12]. Unfortunately, the role of the fibrotic matrix is neglected in the study related to the pathophysiology of muscular dystrophies.

Animal models are generally used to unravel the pathogenesis and test new drugs but their relevance is limited for muscular dystrophies due to deep differences between animals and humans. In fact, these pathological animal models do not accurately recapitulate the severe disease phenotypes observed in human [13]. 2D cellular models relying on the culture of muscle cells on plastic are popular to study muscular dystrophies thanks to their high reproducibility and their easy analysis. Monolayers of myoblasts are cultured on a petri dish coated with Laminin, fibrin or Matrigel® [14–16]. However, the absence of 3D cell organization and cell/ECM interactions do not reflect the physiological condition. Current 3D in vitro muscle models aim to mimic the healthy muscle ECM. The most popular system is based on the encapsulation of muscle cells within hydrogels maintained under tension between two pillars [17,18]. In this system, cells contract hydrogels, facilitating cell alignment and organization. However, mechanical properties of the muscle ECM are not reproduced, nor the biochemical nature of the muscle ECM because this system often consists of fibrin and not dense collagen I [18]. Decellularized ECM has also been employed to mimic the natural environment of skeletal muscle but the mechanical properties of these generated materials seems to be far from the authentic ECM [17]. Beyond their inadequate physical properties, these hydrogels are not porous. As a result, cells located at the center suffer from hypoxia and eventually die, which, in turn, mimics certain pathological conditions [20]. Several attempts have been performed to improve matrices. Bioactive matrices associating muscle and endothelial cells have been designed to create vascularization around muscle fibers [21]. Besides, macroporous hydrogels using a sacrificial matrix made of gelatin have been synthesized to create channels to enhance O_2_ and nutrient diffusion [15]. In these systems, the fibrotic features of the pathological matrix are not reproduced. To the best of our knowledge, the only system focusing on the pathological conditions associate muscle fibers with fibrotic fibroblasts immortalized from patients affected by the Duchenne muscular dystrophy. In this case, the authors only focus on the pathological cell phenotype but the impact of fibrotic matrix is not analyzed [22].

In our group, we have previously developed dense collagen hydrogels mimicking the structure and the physical characteristics of healthy muscle extracellular matrix [23]. The unidirectional 3D printing of dense collagen filaments and their rapid gelling generated macroporous and anisotropic constructs possessing physiological mechanical properties (more specifically E=12 kPa). Myoblasts (C2C12) cultured within large matrix pores dedicated to cell colonization (600 µm in diameter), differentiated into mature myotubes and adopted a 3D organotypic organization mimicking the fascicles of myofibrils [23].

In this study, we aimed to develop a novel 3D in vitro model reproducing the structure and the physical properties of the persistent fibrotic ECM observed in muscular dystrophies. For this purpose, the 3D printing technique was adapted to produce non-porous collagen matrices with high stiffness (40-60 kPa) and altered anisotropy. Then, C2C12 myoblasts were cultured within fibrotic or healthy matrices previously developed. The impact of the fibrotic matrix on C2C12 cell behavior was analyzed at the gene expression level to understand its role in the pathophysiology of muscular dystrophies. Lastly, matrices possessing a single characteristic of fibrotic matrices (i.e high stiffness or non-porosity) were generated to evaluate the impact and predominance of each physical property of the pathological matrix on the cell phenotype.

## 2. Materials and Methods

### 2.1. Solutions preparation

5X PBS solution was prepared with 680 mM NaCl, 40 mM Na2HPO4, 12 H2O, 13 mM KCl and 8 mM NaH2PO4,1 H2O (all products were from Sigma). 1X PBS was obtained by diluting 5 times 5X PBS in distilled water. Ammonium hydroxide 30 % (v/v) (Carlo Erba) was poured in a beaker inside a dessicator to generate vapors.

### 2.2. Collagen extraction and purification

As previously described, type I collagen was extracted and purified from rat tail tendons [23–25]. Briefly, rat tails were rinsed with 70% ethanol and cut into small pieces of 1 cm to extract tendons. After several rinses in 1X PBS, tendons were solubilized in 500 mM acetic acid (Carlo Erba). Then, collagen was precipitated by adding a 4M NaCl (Sigma) solution to obtain a final concentration at 0.7M.

The following day, centrifugations and dialysis were performed to generate a purified collagen solution. Collagen purity was determined after SDS-PAGE electrophoresis (MiniProtean TGX, Biorad) and its concentration estimated by hydroxyproline titration [26]. After an evaporation stage in a safety cabinet for several days, the collagen solution was concentrated up to 30 mg·mL^-1^ (3 % w/v). Finally, the collagen solution was stored at 4°C prior to utilization.

### 2.3. Synthesis of healthy and fibrotic matrices

3D printing was performed using a home-built 3D printer [23]. The concentrated collagen solution was poured into a 1 mL syringe (Terumo) with a flat bottom 23G needle (inner diameter 330 µm). An unique layer of 10 mm x 5 mm x 0.33 mm was designed using AutoDesk Fusion 360 software. The 3D file (.stl) was then sliced with Repetier software with a rectilinear pattern to obtain unidirectional lines inside the square. This layer was printed out and repeated at different z; 5 times to form a 5-layered collagen construct with all filaments oriented in the same direction. The extrusion speed was optimal at 2 mm·s^-1^ (extrusion rate 1 µl·s^-1^). The filling was set to 100% to ensure cohesiveness between every filament but the interlayer gap was tuned to 300 µm to generate intrinsic porosity. Regarding the healthy matrices, anisotropy was induced by collagen shearing during extrusion, and maintained by printing into a 5X PBS bath that induce a fast gelling of collagen filaments (Figure 1). Furthermore, printing in 5X PBS allowed shape retention of collagen filaments and avoided any shrinkage or swelling, For the fibrotic matrices, 3D printing was performed in air with the same printing parameters and gelling was slowly initiated using ammonia vapors. The printing process lasted less than 10 minutes (1 min 30 sec per layer) (Figure 1). An extend collagen gelling was performed after 3D printing on healthy matrices in order to maintain anisotropy and set high mechanical properties. After the initial printing step inside 5X PBS, hydrogels were left for 1 day in a large volume of 5X PBS to continue to gel at pH 7.4 with high ionic strength. At the end of this period, collagen constructs were exposed to ammonia vapors within a dessicator for 1 day to complete gelation and increase stiffness.

**Figure 1:**
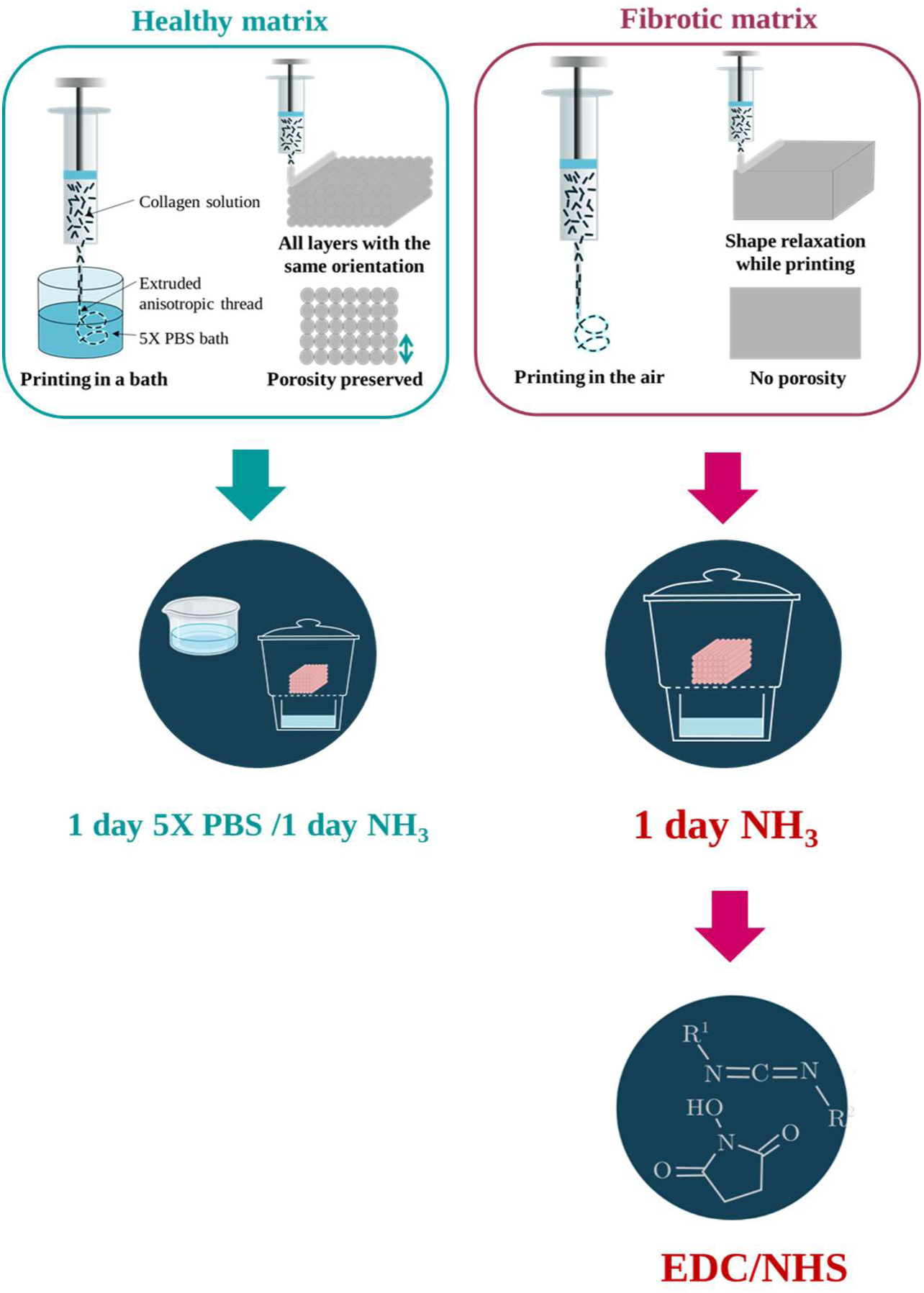
Fabrication of healthy and fibrotic matrices by 3D printing. Healthy matrices were printed in 5X PBS, gelled in 5X PBS for one day and in ammonia vapors for another day. Fibrotic matrices were printed in air, gelled in ammonia vapors for one day and crosslinked using EDC/NHS.

To generate fibrotic matrices, 3D constructs printed in air were only gelled by ammonia vapors (Figure 1). For this purpose, they were placed inside a desiccator for 1 day. Then, all matrices were rinsed several times in 1X PBS until a neutral pH was reached (at least 24 hours).

### 2.4 Cross-linking process to increase the matrix stiffness

Fibrotic matrices required a crosslinking step to increase their mechanical properties. The crosslinker EDC (1-éthyl-3-[3-diméthylaminopropyl] carbodiimide hydrochloride) coupled with NHS (N-hydroxysulfosuccinimide) dissolved in 75% ethanol was used in this study. Tests were first performed on casted hydrogels gelled with ammonia vapors to determine the adequate EDC/NHS concentration to generate fibrotic matrices. For this purpose, different concentrations of cross-linkers were used (Supplementary information - Table 1). Then, cross-linkers at the chosen concentration were used on 3D printed matrices to evaluate their impact on the mechanical properties on 3D constructs. Then, fibrotic matrices were rinsed three times in 1X PBS to remove urea produced by the reaction and the remaining crosslinking molecules. Matrices without any crosslinking or only incubated with 75% ethanol (crosslinking solvent) were used as controls.

**Table 1:** Young’s modulus measured in collagen matrices according to the axis of stretching and determination of anisotropy ratio.

|  |  | Healthy matrix |  | Fibrotic matrix |  |
| --- | --- | --- | --- | --- | --- |
|  |  | 1 day PBS/1 day NH3 |  | 1 day NH3 w/ EDC/NHS |  |
|  |  | E (kPa) | Std (kPa) | E (kPa) | Std (kPa) |
| Toe moduli | Casted | 10,1 | 1,8 | 54,8 | 5,2 |
|  | Printed & stretched longitudinally | 14,9 | 4,8 | 57,5 | 20,5 |
|  | Printed & stretched transversally | 9,5 | 2,4 | 65,0 | 28,3 |
|  | Anisotropy ratio | 1,6 |  | 0,9 |  |
| Linear moduli | Casted | 69,8 | 2,5 | 286,1 | 18,5 |
|  | Printed & stretched longitudinally | 88,0 | 14,4 | 334,4 | 146,0 |
|  | Printed & stretched transversally | 57,1 | 9,3 | 357,2 | 86,8 |
|  | Aniotropy ratio | 1,5 |  | 0,9 |  |

### 2.5 Synthesis of matrices possessing a single characteristic of fibrotic matrices

To evaluate the impact of stiffness or porosity on cell phenotype, two matrices were designed. A rigid and porous matrix was developed from healthy matrices. After producing a healthy matrix, the cross-linking step used for fibrous matrices was applied to healthy matrices to increase their rigidity (Supplementary Information 1). Additionally, soft, non-porous matrices were designed using the same process as for fibrous matrices but without the cross-linking step (Supporting Information 1).

### 2.6 X-Ray Microtomography (Micro-CT) analysis

Micro-CT was performed to monitor porosity within the different matrices. Matrices were incubated with Micropaque contrast agent (Guerbet) before their observation using a high-resolution X-ray micro-CT system (Quantum FX Caliper, Life Sciences, Perkin Elmer) hosted by the PIV Platform (UR2496). The contrast agent appeared white and collagen filaments grey. Standard acquisition settings were applied (voltage 90 kV, intensity 160 mA), and scans were performed with a field of view of 1 cm^2^. Micro-CT datasets were analyzed using the built-in multiplanar reconstruction tool, Osirix Lite (Pixmeo), to obtain time series of images and 3D reconstruction. Micro-CT images were thus analyzed using ImageJ. Images were thresholded and the diameter of the pores was measured. The irregular shape of the pores was captured using the following parameters: Feret max diameter (the smallest circle that surrounds the pore) and circularity.

### 2.6. Polarized Light Optical Microscopy (PLOM)

Polarized Light Optical Microscopy was used to observe anisotropy in matrices. PLOM was performed using a photonic Nixon microscope equipped with crossed-polarizers to observe collagen birefringence. Collagen fibrils with uniaxial alignment will increase or decrease the light intensity depending on the relative orientation of the crossed polarizer. A series of images were acquired with multiple sample orientations to demonstrate the light intensity variation. Maxima and minima were separated by 45°.

### 2.7 Second-harmonic generation (SHG)

Second-harmonic generation microscopy was used to observe anisotropy as a strong SHG signal is proportional to collagen quantity and alignment. SHG images were acquired with a Mai Tai multiphoton laser-equipped confocal microscope (Leica SP8). SHG signal was collected with a hybrid detector between 430 and 450 nm (excitation at 880 nm). A long-distance objective (25X water immersion) was used to acquire z-stacks with optimal settings for each sample. Image series were acquired with the Leica Application Suite X software.

### 2.8 Rheological measurements

Shear oscillatory measurements were performed on casted collagen hydrogels with different gelling parameters using an Anton Paar rheometer. An 8 mm parallel-plate geometry was fitted with a rough surface to avoid gel slipping. All measurements were performed at 37 °C. Storage modulus, G’ and loss modulus, G” were recorded during a frequency sweep from 0.1 to 10 Hz with an imposed strain of 1%. This strain corresponds to non-destructive conditions (linear viscoelastic regime) as previously checked (data not shown). The geometry and a stabilized normal force (0.01 N) were optimized. Three samples of each matrix were tested.

### 2.9 Tensile tests

Mechanical tests were performed on printed gels with a tensile testing machine (Instron) equipped with a 10 N load cell and Bluehill software. Printed gels were obtained by printing five successive layers of 25 mm x 5 mm either in the longitudinal or transversal direction. Total thickness of the sample was around 1 mm (measured before each experiment). All experiments were performed on hydrated samples (three specimens per condition) at 21 °C at a constant strain rate of 0.06 s^-1^. The elastic (Young’s) modulus of collagen hydrogels was measured by a linear fit within the first 5 % deformation (Toe Young’s modulus). The strain, “e” and stress, “s” were estimated as e=ΔL/L (L=sample length) and s=F/S0 respectively. Cross-section area, S0, was assumed to be homogeneous and calculated from the measure of the initial width and thickness.

### 2.10 Crosslinking cytotoxicity

Normal Human Dermal Fibroblasts (NHDF) were used to measure the potential cytotoxicity of crosslinking. Briefly, cells were cultured in a complete cell culture medium (Dulbecco’s Modified Eagle’s Medium (DMEM) supplemented with 10 % (v/v) fetal bovine serum, 100 U·mL^-1^ penicillin, 100 μg·mL^-1^ streptomycin, 0.25 μg·mL^-1^ Fungizone, and Glutamax) for 48 hours. Then, they were trypsinized (with 0.1 % (v/v) trypsin and 0.02 % (v/v) EDTA), and 5,000 cells were seeded on top of crosslinked collagen hydrogels and cultivated for 24 hours. Then metabolic activity was measured by an Alarm blue test and a live/dead assay as previously described [27].

### 2.11 Generation of large channels dedicated to cell culture

Large channels were generated within collagen matrices using needles. These channels were created to culture myoblasts and allow their organization in fascicles. First, the matrices were printed into a 5X PBS bath or in air, but inside dedicated 3D-printed plastic molds with holes (10 x 5 x 1.5 mm) (Supplementary Information 2A). After 30 min, 2 needles (external diameter 600 µm/23 G) were placed into the collagen constructs through the dedicated holes (Supplementary Information 2A). The fibrillogenesis and gelling in 5X PBS (healthy or stiff, porous matrices) or NH_3_ (fibrotic or soft, non-porous matrices) were continued over 24H. After complete gelling with ammonia vapors (healthy or stiff, porous matrices) or cross-linking with EDC/NHS (fibrotic or soft, non-porous matrices), the constructs were rinsed in 1X PBS and needles removed to reveal straight, well-defined channels that cross the whole hydrogel.

### 2.12. C2C12 culture inside large channels

C2C12 murine myoblasts were expanded in proliferation medium (DMEM high glucose, 20 % (v/v) fetal bovine serum, 100 U·mL^-1^ penicillin, 100 μg·mL^-1^ streptomycin). To seed C2C12 myoblasts into the different matrices, cells were resuspended in Matrigel® at 3·10^7^ cells.mL^-1^ (Sigma, Bioreagent) on ice. 3 µl of cell suspension were injected with a P10 pipet in each channel created by needles (600 µm). After 24h in proliferation medium, cells were cultured in differentiation medium (DMEM high glucose, 2 % (v/v) horse serum, 100 U·mL^-1^ penicillin, 100 μg·mL^-1^ streptomycin) for 5 days. The medium was changed every day. Constructs were then fixed in 4 % paraformaldehyde (w/v) (Sigma) in 1X PBS overnight at 4 °C and sectioned into 250 µm slices with a vibratome. All experiments were carried out in triplicates.

### 2.13 Immunofluorescence

For healthy and fibrotic matrices colonized by C2C12, fluorescent labeling of nuclei (TOPRO-3, Thermofisher) and F-actin filaments (Alexa Fluor 488 phalloidin, Thermofisher) were performed on 250 µm sections prepared using vibratome. Additional labeling of C2C12 with MF-20 hybridoma mouse IgG2B primary antibody and Alexa Fluor 546 goat anti-mouse IgG2B secondary antibody (Invitrogen) was performed to evaluate the cell differentiation into myotubes. Observations were conducted with a Leica SP5 upright confocal, multiphoton laser scanning microscopy, which enabled the simultaneous acquisition of fluorescence and second-harmonic generation signals.

### 2.14 RNA sequencing

Matrices were mixed with 1 mL of Trizol reagent ® (Invitrogen) and homogenized using an Ultraturax. For 30 second grinding periods were alternating with 30 sec on ice. After a centrifugation step at 10000 g, for 5 min, the supernatants were collected and mixed with 100 μL of chloroform for 15 minutes. A new centrifugation (10 min, 10,000 g, RT) was carried out to isolate the aqueous fraction. Then, RNA was extracted and purified following the guidelines of QIAGEN Minikit (QIAGEN). RNA samples were stored at −80°C prior to utilization. Six healthy and 6 fibrotic matrices were processed.

After RNA extraction, RNA concentrations were obtained using nanodrop or a fluorometric Qubit RNA assay (Life Technologies, Grand Island, New York, USA). The quality of the RNA (RNA integrity number) was determined on the Agilent 2100 Bioanalyzer (Agilent Technologies, Palo Alto, CA, USA) as per the manufacturer’s instructions.

To construct the libraries, 200 ng of high quality total RNA sample (RIN >8.2) was processed using Stranded mRNA Prep kit (Illumina) according to manufacturer instructions. Briefly, after purification of poly-A containing mRNA molecules, mRNA molecules are fragmented and reverse-transcribed using random primers. Replacement of dTTP by dUTP during the second strand synthesis will permit to achieve the strand specificity. Addition of a single A base to the cDNA is followed by ligation of Illumina adapters. Libraries were quantified by Q bit and profiles were assessed using the DNA High Sensitivity LabChip kit on an Agilent Bioanalyzer. Libraries were sequenced on an Illumina Nextseq 500 instrument using 75 base-lengths read V2 chemistry in a paired-end mode.

After sequencing, a primary analysis based on AOZAN software (ENS, Paris) was applied to demultiplex and control the quality of the raw data (based of FastQC modules / version 0.11.5). Obtained fastq files were then aligned using STAR algorithm (version 2.5.2b) and quality control of the alignment realized with Picard tools (version 2.8.1). Reads were then counted using Featurecount (version Rsubread 1.24.1) and the statistical analyses on the read counts were performed with the DESeq2 package version 1.14.1 to determine the proportion of differentially expressed genes between two conditions.

### 2.15 RNA-Seq data analysis

Analysis of sequencing data quality, reads repartition (*e.g.*, for potential ribosomal contamination), inner distance size estimation, genebody coverage, strand-specificity of library were performed using FastQC v0.11.2, Picard-Tools v1.119, Samtools v1.0, and RSeQC v2.3.9. Reads were mapped using STAR v2.4.0f1 [28] on the mouse mm39 genome assembly and read count was performed using featureCount from SubRead v1.5.0 and the Mouse FAST DB v2022_1 annotations.

Gene expression was estimated as described previously [29]. Only genes expressed in at least one of the two compared conditions were analyzed further. Genes were considered as expressed if their FPKM value was greater than FPKM of 98% of the intergenic regions (background). Analysis at the gene level was performed using DESeq2 [30]. Genes were considered differentially expressed for fold-changes ≥1.5 and p-values ≤0.05.

Pathway enrichment analyses and GSEA analysis on MSigDB hallmark gene sets were performed using WebGestalt v0.4.4 [31] merging results from up-regulated and down-regulated genes only, as well as all regulated genes. Pathways and networks were considered significant with p-values ≤0.05.

### 2.16 Statistical Analysis

All experiments were carried out at least twice, and the results were expressed as the mean values ± standard deviation (SD). The differences were analyzed using Kruskal-Wallis test followed by Dunn’s test for multiple comparisons. *p* < 0.05 was considered significant.

## 3. Results

In this study, the synthesis of porous collagen hydrogels previously developed in our group was adapted to design fibrotic matrices. First, the intrinsic porosity and anisotropy generated by the 3D printing in healthy matrices was annihilated by modifying the gelling conditions. Then, none porous matrices were cross-linked by different concentrations of EDC/NHS to increase their Young’s modulus up 50 kPa and decrease their deformability. Then, the behavior of C2C12 myoblasts was studied in healthy and fibrotic collagen behavior to assess the impact of fibrosis on muscle regeneration and phenotype. Finally, the cell phenotype in soft, non-porous or stiff, porous matrices was compared to that in healthy matrices to determine which pathological characteristic (stiffness or low porosity) was predominant for the cellular phenotype in fibrotic matrices.

### 3.1 Intrinsic macroporosity within healthy and fibrotic matrices

The intrinsic porosity generated by 3D printing was analyzed in healthy and pathological matrices. Healthy matrices, printed in a 5X PBS solution, exhibited elongated channels crossing the construct (Figure 1 and 2A). This was due to the stacking of collagen filaments, their shape retention and their rapid gelling in PBS 5X in solution. In contrast, the shape relaxation of collagen occurred in pathological matrices due to the slow gelling by ammonia vapors. As a consequence, filaments lost their cylindrical shape, spread and filled the pores (Figure 1). After gelling, a non-porous construct was generated (Figure 2A and C). Pores analysis confirmed the absence of pores in the fibrotic matrix whereas a minimal Feret diameter of 100 µm in average with 10 pores per slice was detected for the healthy matrix (Figure 2B and C).

**Figure 2:**
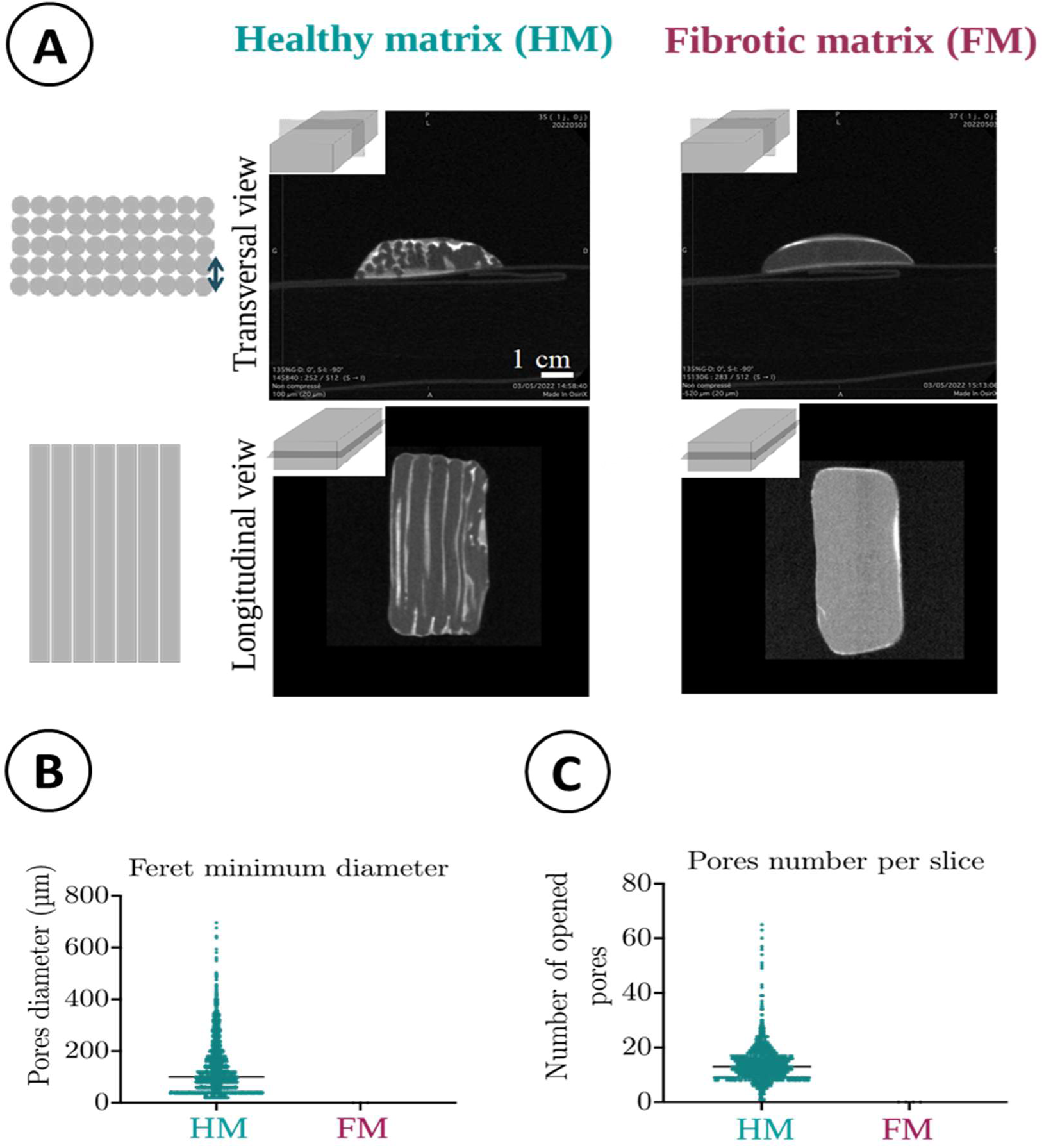
Intrinsic porosity of healthy and fibrotic matrices. (A) Macroporosity observed by X ray-microtomography. (B) Pore size determined by measurement of the Feret minimum diameter. (C) Number of pores on each hydrogel section (n=4). Black bar: median.

### 3.2 Tuning the mechanical properties of fibrotic matrices

Compared to healthy matrices, fibrotic matrices are at least three times stiffer (E = 30-75 kPa). To mimic the mechanical properties of fibrotic matrices, different concentrations of EDC/NHS were tested to crosslink and augment matrix rigidity. This crosslinking step was first performed on casted hydrogels after complete collagen gelling with ammonia vapors. Increasing the concentration of EDC/NHS in casted matrices had a strong impact on their rheological properties. Incubating matrices for 2 hours with an EDC/NHS solution concentrated at 5/1.2 and 20/4.8 mg.mL^-1^ increased the storage modulus up to c.a 15 and 25 kPa, respectively (Figure 3). Interestingly, 75% ethanol, the solvent of EDC/NHS, did not significantly impact collagen hydrogel mechanical properties (Figure 3). The two red lines symbolize the stiffness range corresponding to pathological and fibrotic extracellular matrices. Healthy and fibrotic matrices are highly hydrated collagen hydrogels, with typical 97 wt% of aqueous solution. It means that the matrix can be assumed to be nearly incompressible and the Young’s modulus, E can reasonably be estimated around ≈ 3G’. Hence, E is approximated 45 kPa and 75 kPa when matrices are gelled with NH_3_ and cross-linked with 5/1.2 and 20/4.8 mg.mL^-1^ EDC/NHS, respectively. These Young’s modulus values are similar to those measured in pathological muscular matrices (30-70 kPa).

**Figure 3:**
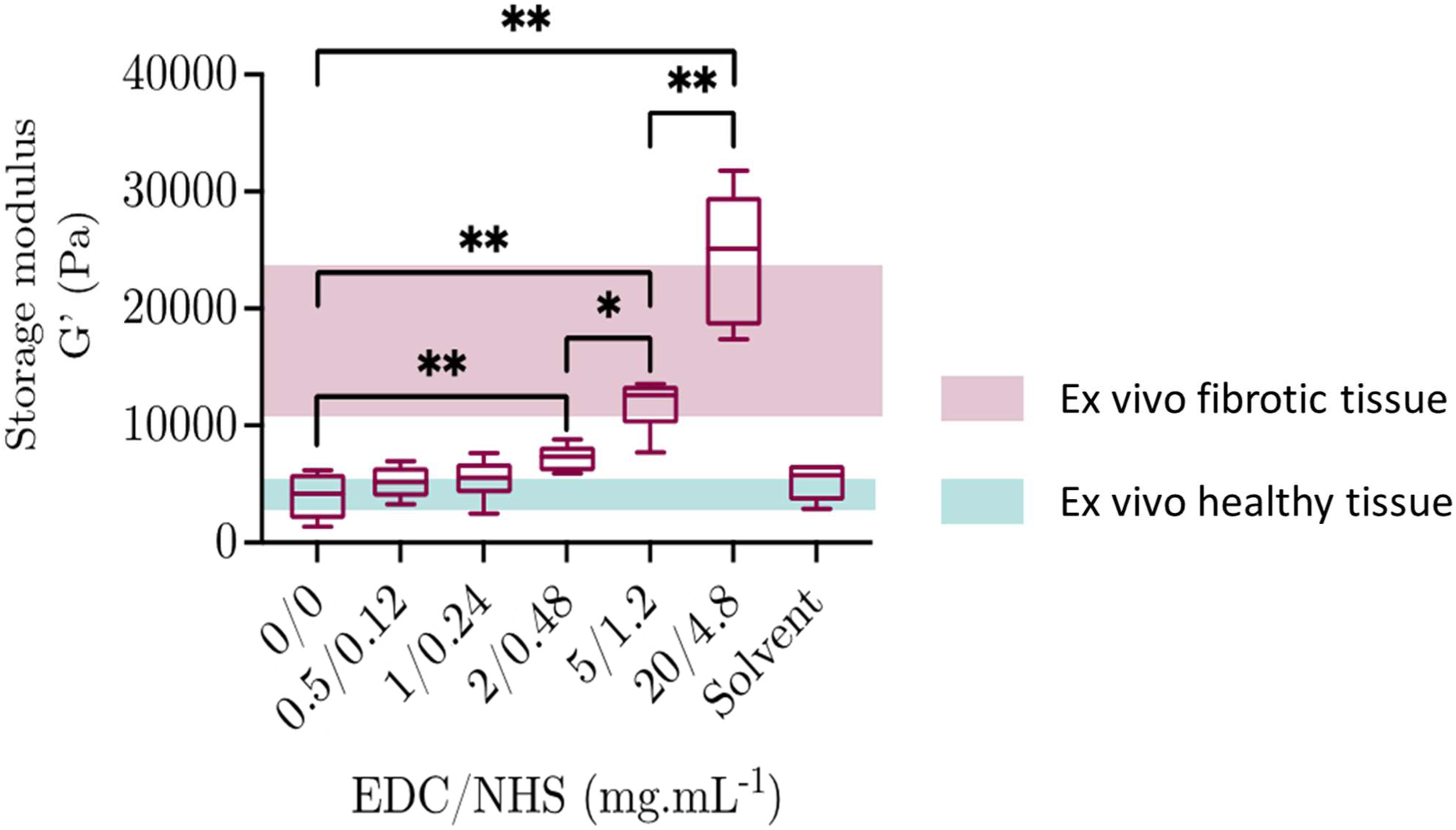
Rheological properties of casted collagen matrices after crosslinking with EDC/NHS. The red and green stripe symbolizes the stiffness range corresponding to fibrotic and healthy extracellular matrices, respectively (n=4); *: *p* ≤0.05.

The effect of the EDC/NHS on the casted collagen hydrogels was then evaluated over time with EDC/NHS=5/1.2 mg·mL^-1^ (Supplementary Information 3). The G’ values reached a plateau between 20 and 25 kPa after 6 hours of reaction (Supplementary Information 3). The loss modulus G” also reached a plateau after 6 hours (data not shown). These results showed that collagen hydrogels cross-linked with EDC/NHS concentrated at 5/1.2 mg·mL^-1^ for 6 hours, exhibit a stiffness similar to the fibrotic muscular extracellular matrix.

### 3.3 EDC/NHS cytotoxicity

EDC/NHS was chosen for its cytocompatibility, but this aspect must be checked before cellularization of the healthy and fibrotic matrices. To do so, normal human dermal fibroblasts were seeded on top of cross-linked hydrogels and their metabolic activity is assessed by Alamar blue assay (Supplementary Information 4A). Cell metabolic activity was only impacted when the highest cross-linkers concentration was used (EDC/NHS= 20/5 mg·mL^-1^). To confirm these results a Live/Dead assay was performed, none of the tested conditions was cytotoxic (Supplementary Information 4B). Hence, the cross-linking using EDC/NHS=5/1.2 mg·mL^-1^ for 6 hours seem not to affect cell survival and was chosen for the cross-linking of 3D printed pathological matrices.

### 3.4 Characteristics of the fibrotic matrices

One characteristic of the pathological matrix is the anisotropy disturbance. To observe intrinsic collagen alignment within matrices, Polarized Light Microscopy (PLM) was first used. When printed in PBS, collagen filaments were anisotropic whereas they loosed their molecule alignment when printed in air due to the collagen relaxation before gelling. Only healthy matrices were birefringent (Figure 4). These results were confirmed using Second Harmonic Generation microscopy (SHG) that showed the low SHG signal in fibrotic matrices (Figure 4).

**Figure 4:**
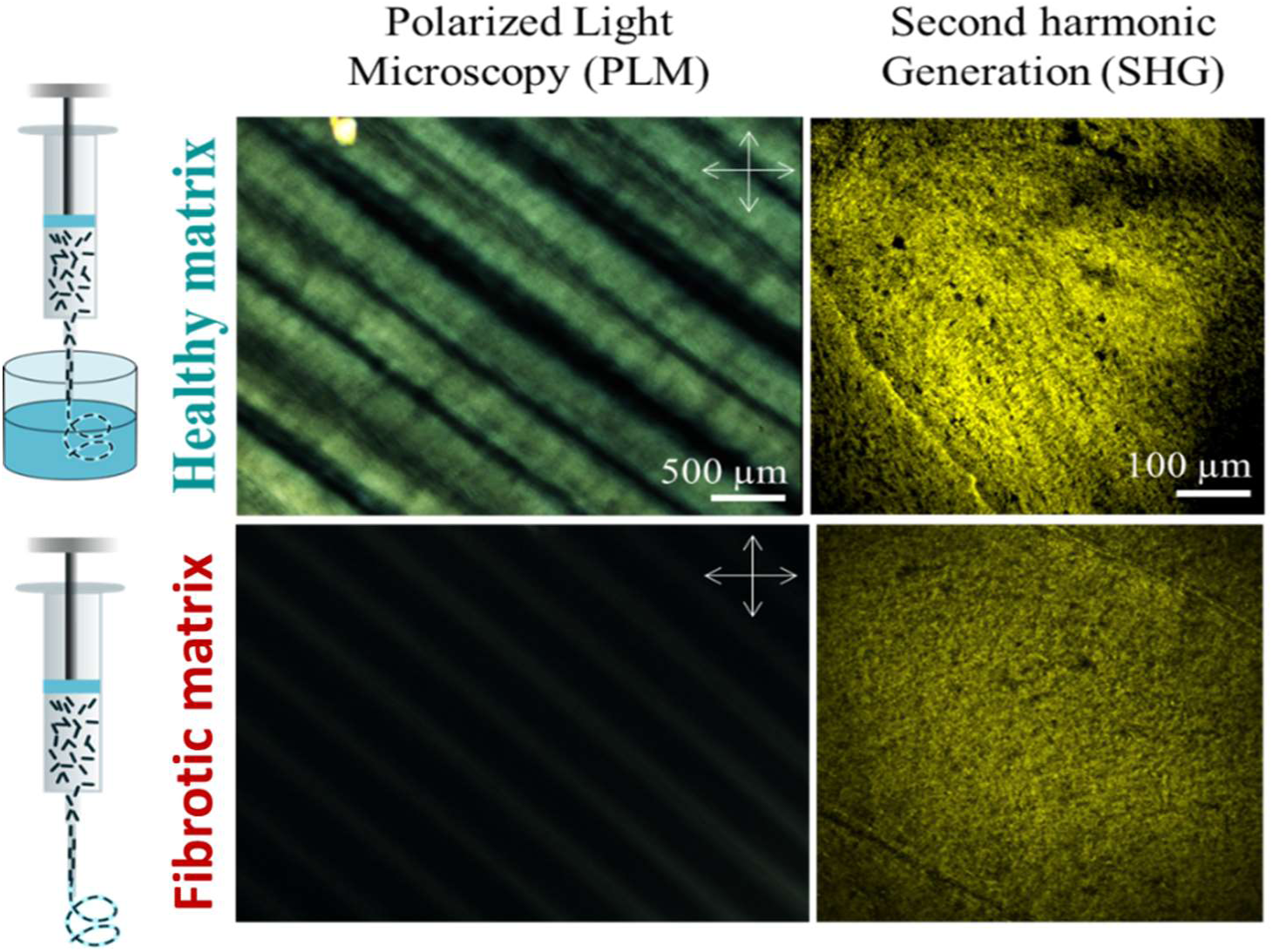
Anisotropy of healthy and fibrotic collagen matrices observed by polarized light microscopy and second harmonic generation microscopy. The absence of birefringence in fibrotic matrices evidences their isotropic structure

The matrix anisotropy was also assessed by analyzing tensile behavior in parallel and perpendicularly to extrusion direction. (Figure 5). For this purpose, printed hydrogels were stretched along the fiber orientation or perpendicularly (dotted curves). As expected from biological tissues, tensile behavior displayed a J-shaped response with a toe region followed to strain hardening region [32,33]. Tensile toe and linear moduli were respectively determined at 5% and 20% strain. Regarding the healthy matrices, the axis of stretching drastically impacted the mechanical behavior (Figure 5A and Supplementary Information 5A for magnified graph). First, both curves generated by the different stretching methods did not overlay. Second, the toe Young’s modulus, calculated at low deformation, was around 15 kPa (Figure 5C) when matrices were longitudinally stretched whereas it decreased down to c.a 9 kPa when hydrogels were perpendicularly stretched (Table 1). The anisotropy ratio, being the ratio between longitudinal and perpendicular moduli, was then calculated to obtain 1.6, thereby showing the presence of macroscopic anisotropy. Moreover, the healthy matrix was highly deformable (above 150%) when stretched along the fibril orientation, but less (below 80%) after a perpendicular stretching (Figure 5A).

**Figure 5:**
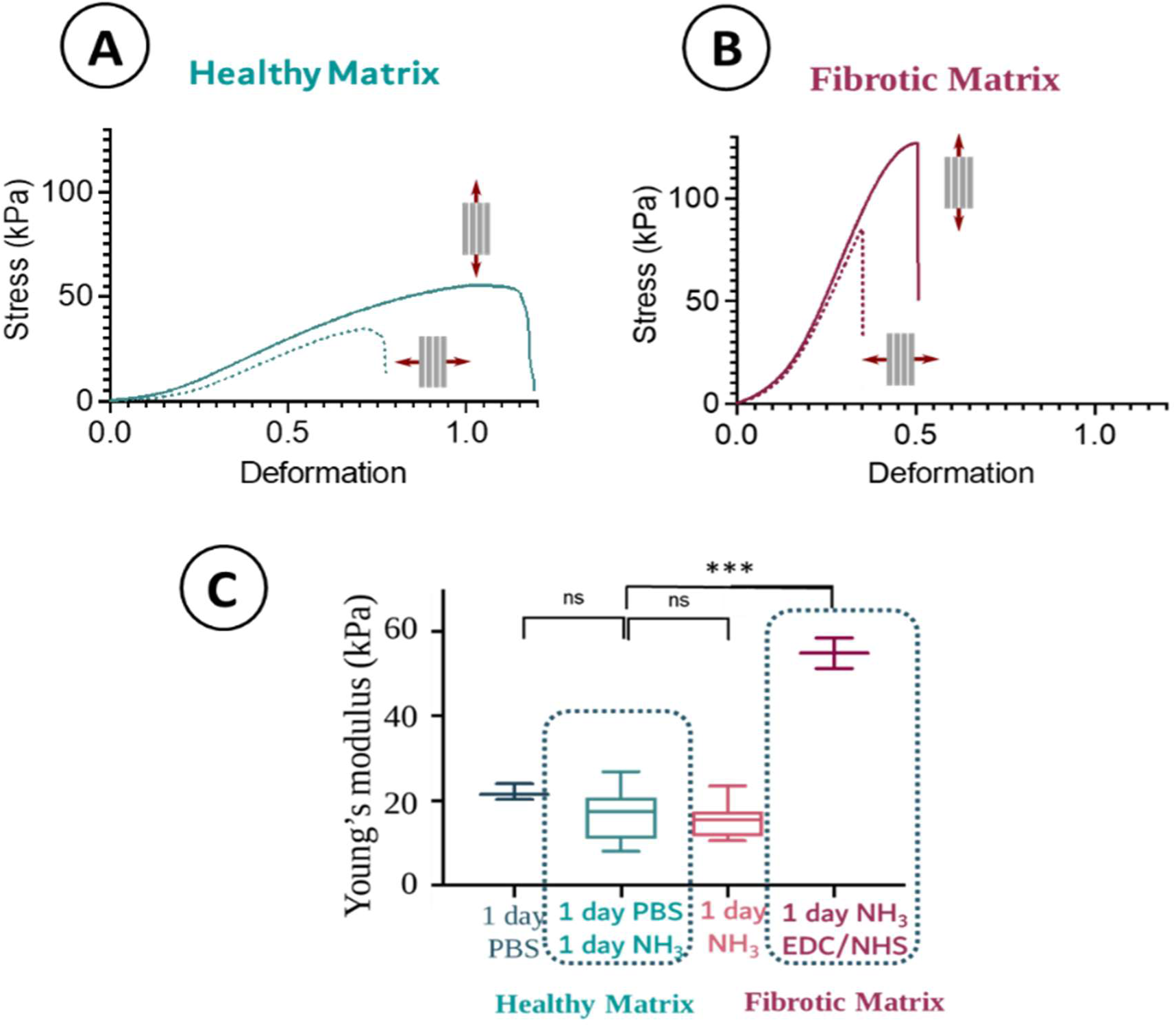
Mechanical properties of 3D printed healthy (A) and pathological matrices (B) assessed by tensile tests. Matrices were longitudinally stretched (plain lines) or perpendicularly stretched (dotted lines). (C) Young’s modulus of dense collagen matrices after of EDC/NHS crosslinking (n=4); *: *p* ≤0.05.

When the fibrotic matrix was analyzed, we noticed that the cross-linking mainly increased the toe and linear moduli and decreased notably the matrix deformability (50%) (Figure 5B). The toe Young’s modulus increased 4 times for cross-linked constructs to reach around 50 kPa (Figure 5C). In addition, the curves obtained from different stretching methods (along fibril orientation and perpendicular) almost overlaid (Supplementary Information 5B), with an anisotropy ratio around 0.9 (Table 1), thereby evidencing the absence of significant anisotropy. Hence, printing in the air affects the macroscopic anisotropic properties of the collagen constructs, whereas the crosslinking step increased its Young’s modulus and reduced deformability.

### 3.5 C2C12 colonization, proliferation and differentiation in healthy or fibrotic matrices

C2C12 were encapsulated within pure Matrigel® and injected into the large channels obtained by the use of 600 µm diameter needles. This size of channels was chosen to allow cells to proliferate, differentiate and adopt a physiological organotypic organization in fascicles characterized by cell alignment and cell/cell contacts in 3D. After 1 day in proliferation medium followed by 5 days in differentiation medium, large mature myotubes expressing myosin heavy chains were observed in healthy matrices (Figure 6). Cells were densely-packed due to the high cell density at seeding and their proliferation within the construct (Figure 6). Moreover, the myotubes were aligned and multinucleated. C2C12 cells had remodeled Matrigel and entirely filled the pores with some cell/cell contact visible (Supplementary Information 6). In contrast, the cell density observed within pores dedicated to cell colonization was lower in fibrotic matrices. Some myotubes were visible, but they were randomly oriented, short, and mononucleated. Some debris were also present.

**Figure 6:**
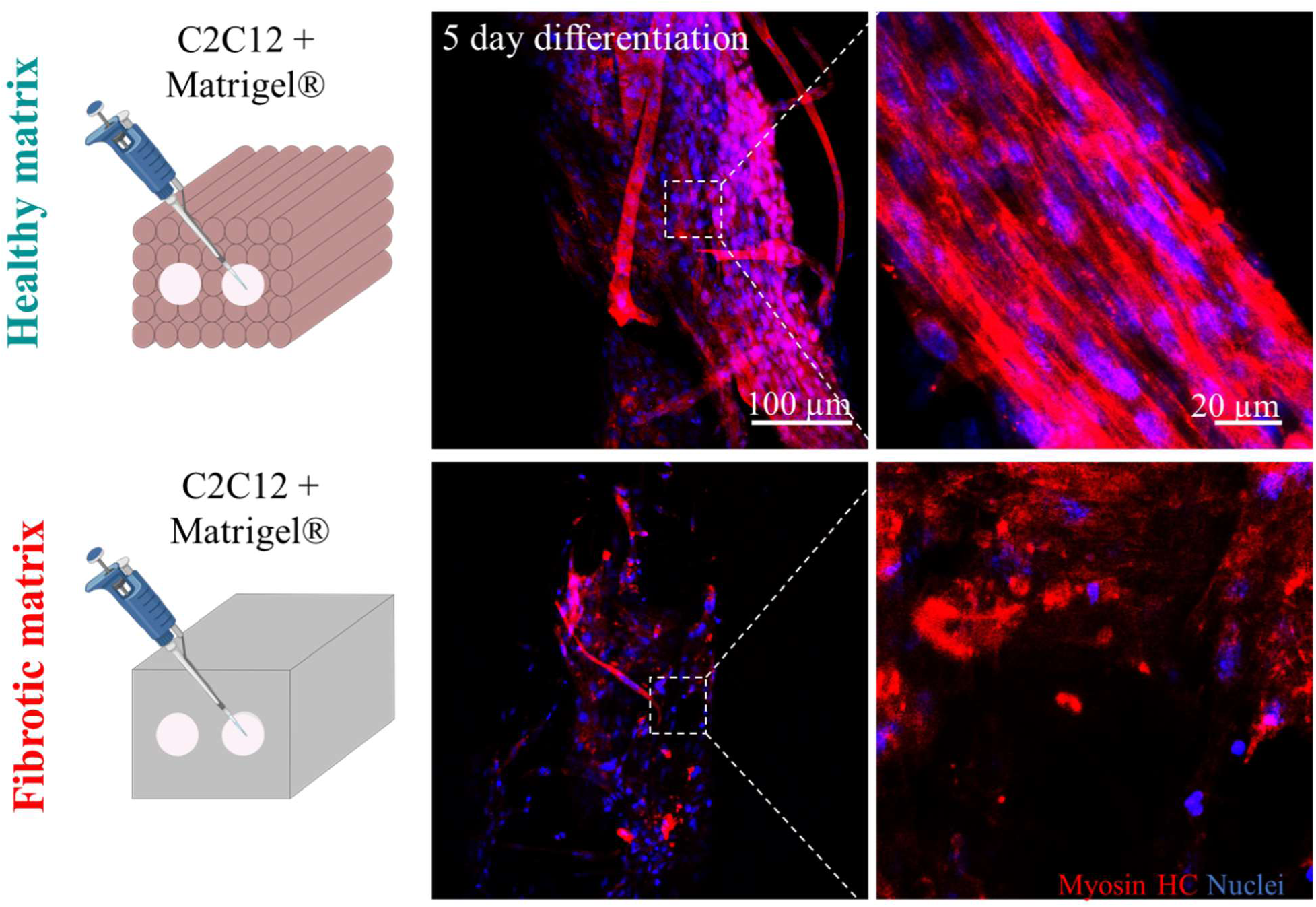
C2C12 differentiation within healthy or fibrotic matrices. C2C12 myoblasts were embedded into Matrigel® and injected within the large channels dedicated to cell colonization; Observation of Myosin Heavy Chain labeling (red) evidenced cell differentiation into mature myotubes in healthy matrices. Nuclei were stained in blue with DAPI.

### 3.6 Gene expression of C2C12 cultivated within healthy or fibrotic matrices

The comparison of gene expression of C2C12 myoblasts cultivated within healthy or fibrotic matrices was performed after 5 days in differentiation medium, when C2C12 cells should be differentiated into mature myotubes. The culture within fibrotic matrices was associated with the regulation of 1652 genes, 888 genes were upregulated and 764 were downregulated (Figure 7). All the data and analysis from RNA sequencing are included in supplementary information files. Then, a Gene Set Enrichment Analysis (GSEA) was performed to determine whether a priori defined sets of genes were statistically modulated and may have an association with the fibrotic matrix.

**Figure 7:**
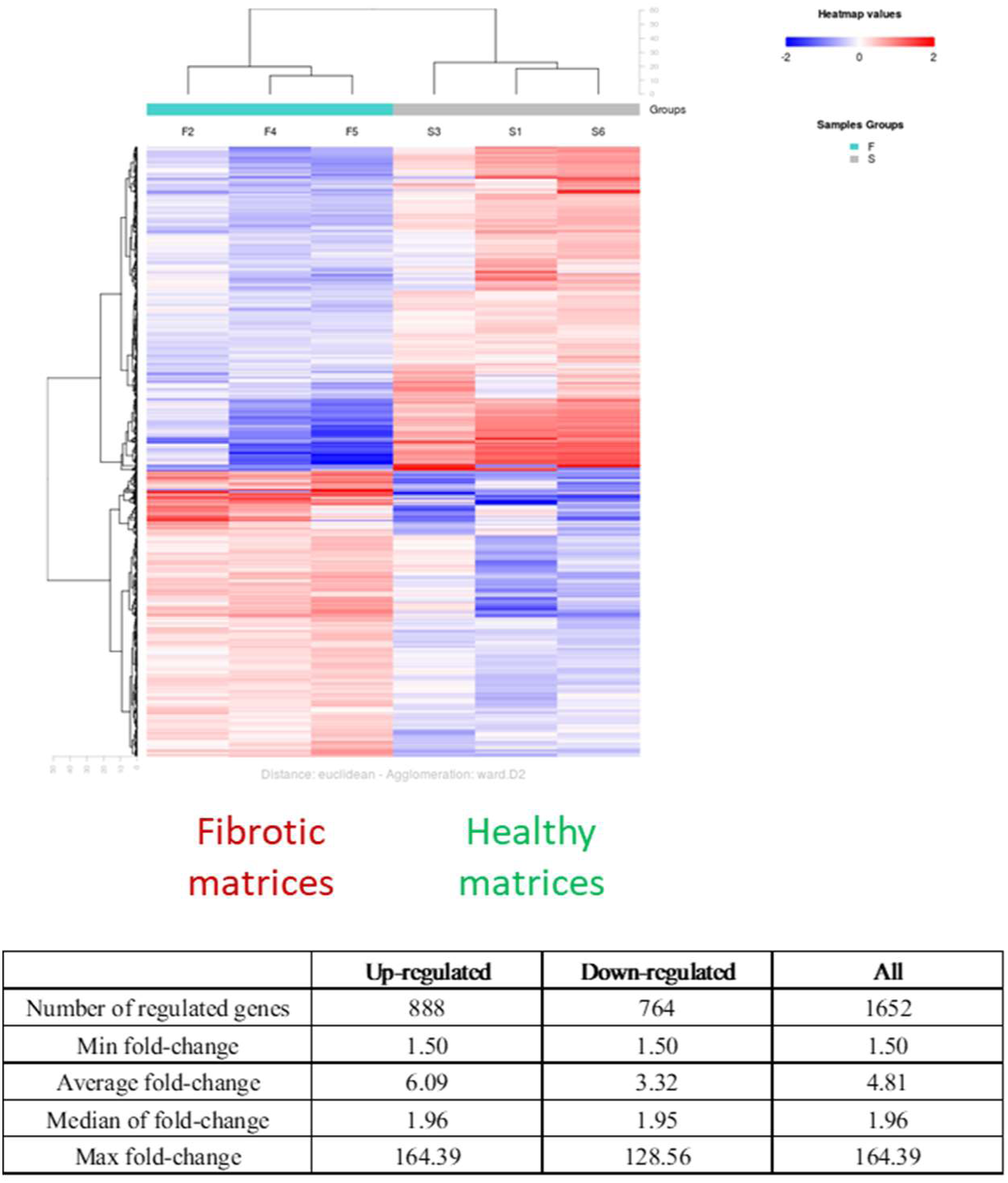
Hierarchical Clustering of Regulated Genes. Comparison of gene expression between C2C12 cultured within healthy and fibrotic matrices. Values taken into account: Fold-change ≥ 1.5 and *p* ≤ 0.05. n = 3 for each group.

#### 3.6.1 Hypoxia

One significant consequence of the absence of porosity in the pathological matrix is the occurrence of hypoxia. The GSEA showed that genes related to the response to hypoxia were upregulated in fibrotic matrices (Figure 8). A leading edge number of 87 was found in a set of 152 genes (See GSEA graphs). To go further, a pathway analysis using KEGG database revealed that genes regulating the synthesis or the degradation of HIF1α (Hypoxia Induced Factor 1 alpha) were upregulated (Link to KEGG pathway: mmu04066). Surprisingly, Egln 1, Egln 3 and vhl, proteins degrading HIF1α in normoxia were upregulated (Supplementary Information 7). IL6 and its receptor that promote HIF1 α synthesis were upregulated. Last, the MAP kinase signaling pathway involved in the synthesis of this factor was also upregulated. The evidence of hypoxic conditions in fibrotic matrices was confirmed by the upregulation of angiogenic factors such as angiopoetin1, VEGFA, VEGFB and VEGFD (Supplementary Information 8). These factors are produced in hypoxic conditions to counterbalance the reduced access to nutrients and oxygen. Another characteristic of hypoxia in muscle is the metabolic switch toward glycolysis. The GSEA revealed that 151 genes involved in the glycolysis were upregulated with 44 as leading edge number (Figure 8). Using the Panther database, 9 genes were upregulated in the glycolysis pathway (link to Panther pathway: Am P00024). Among them, enzymes involved in the intracellular accumulation of glucose such as hexokinases were upregulated. In addition, glycolysis in hypoxia leads to lactate production. In fibrotic matrices, the lactate deshydrogenase upregulation was observed to detoxify the muscle (Supplementary Information 7). Hypoxia and glycolysis leads to the production of reactive oxygen species in vivo. The GSEA showed that 37 genes related to ROS were modulated in fibrotic matrices (Figure 8) with a trend towards upregulation (leading edge number: 17). In addition, antioxidant enzymes such as superoxide dismutase or glutathione peroxidase are also upregulated to decrease the ROS quantity (Supplementary Information 7).

**Figure 8:**
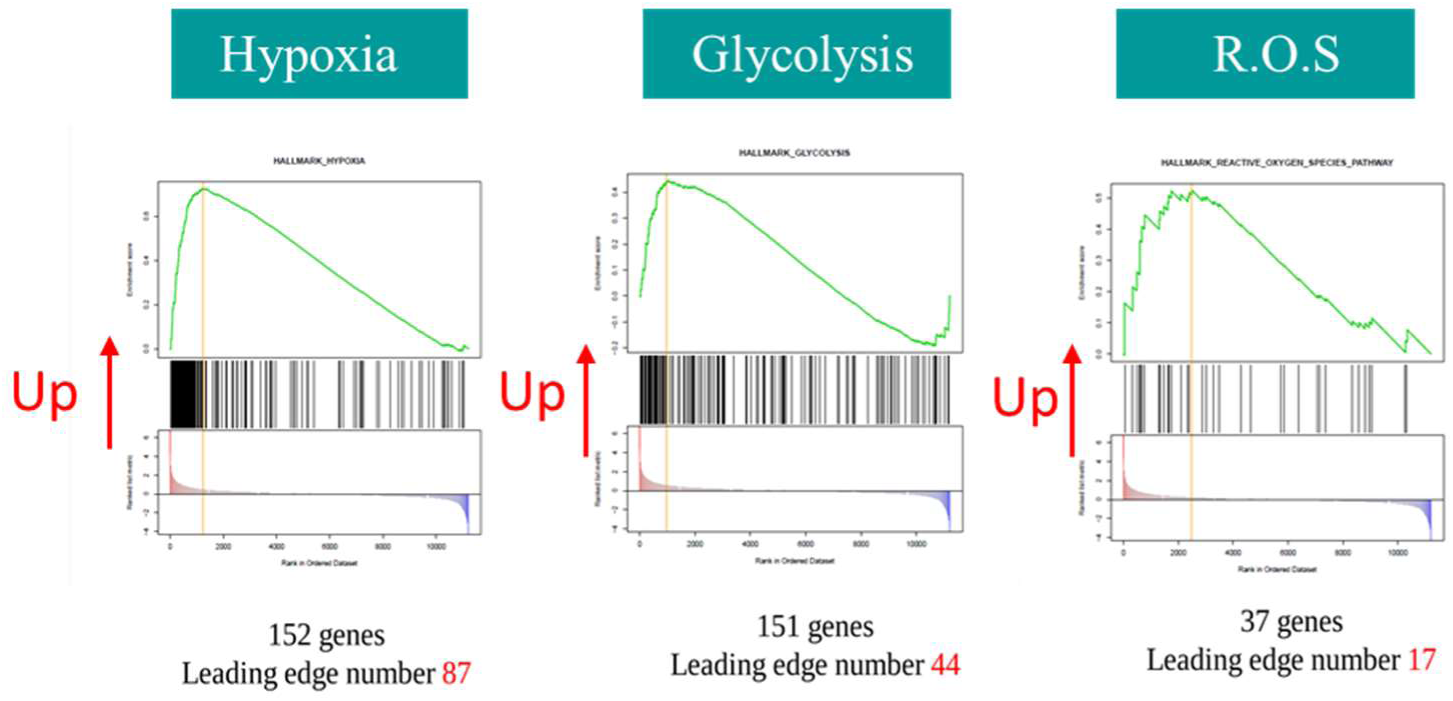
Upregulation of target genes in response to hypoxia shown by GSEA

#### 3.6.2 Inflammation

The fibrotic matrix likely induces a reduced availability of oxygen which in turn generated inflammation. The expression of inflammatory molecules such as TNFα, IL-1 and IL-6 were upregulated in fibrotic matrices. More generally the TNFα signaling via NFϰB pathway and the “inflammatory response” pathways were upregulated (Figure 9).

**Figure 9:**
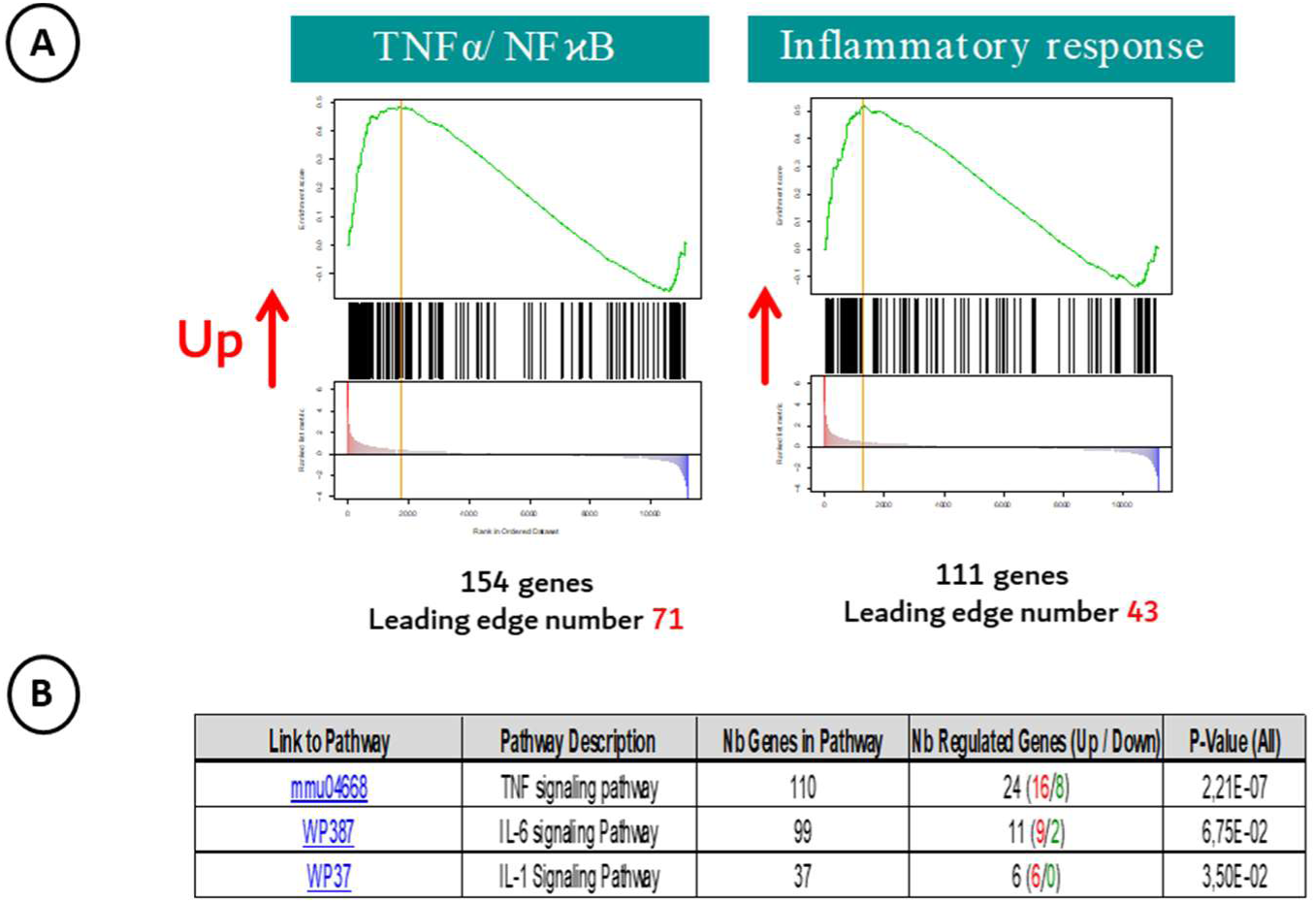
(A) Upregulation of inflammatory response in fibrotic matrices analyzed by GSEA (B) Upregulation of specific cytokines pathways involved in inflammation (Wiki and Kegg database).

#### 3.6.2 Cell cycle

The bioinformatics analysis revealed a significant impact of fibrotic matrices on cell cycle. In pathological matrices, cell cycle was stopped (Figure 10A). The different phases of cell cycle were impacted (G1, S, M) (Figure 10B). In addition, genes involved in the transitions and checkpoints between the different phases were also downregulated (G1/S, G2/M checkpoint). Last, several pathways related to mitotic phases were strongly impacted in the fibrotic matrix (prophase, metaphase, anaphase, etc…) (Figure 10B).

**Figure 10:**
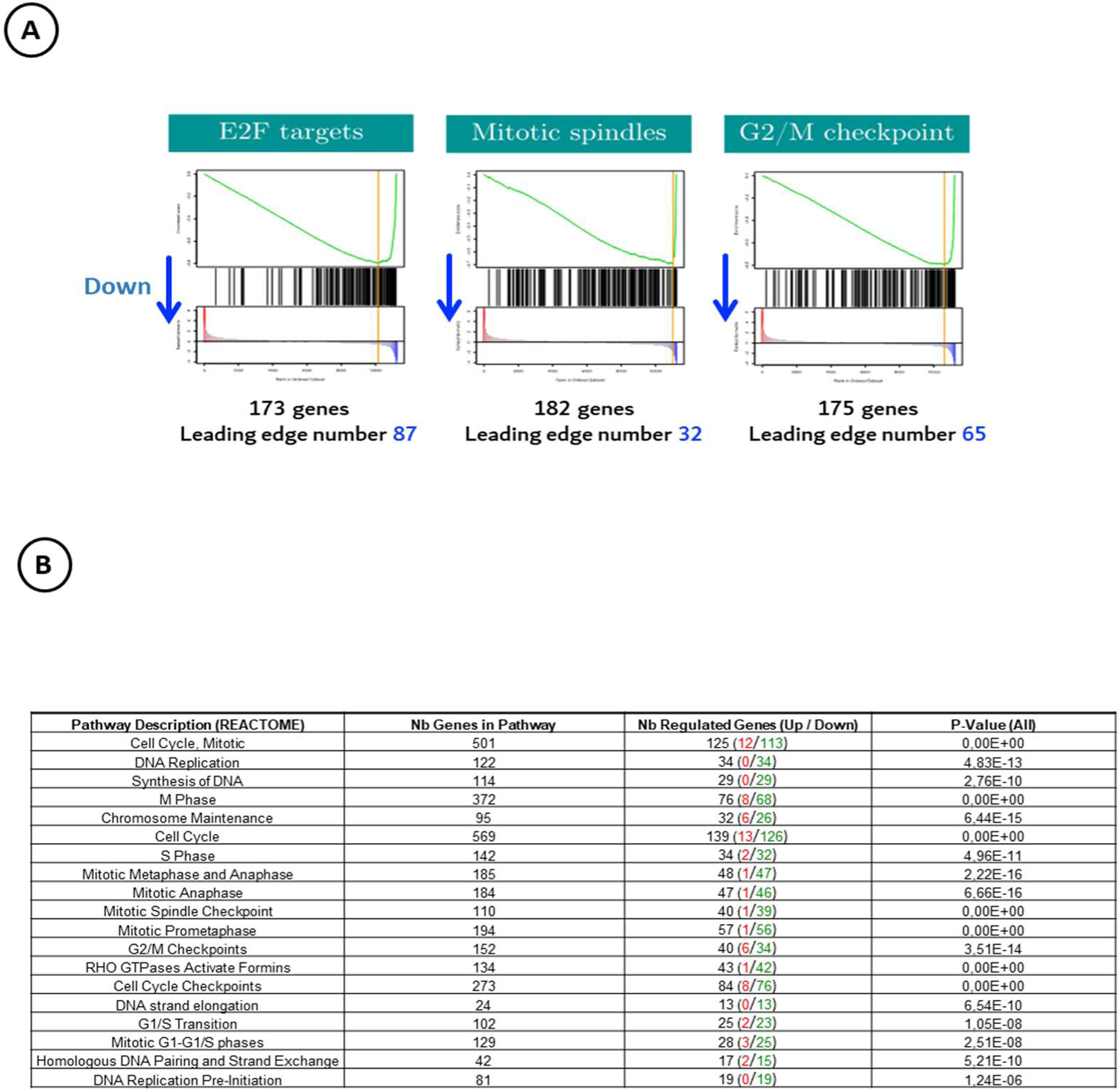
Impact of fibrotic matrix on cell cycle of C2C12 myoblasts. (A) Analysis of cell cycle by GSEA. (B) List of pathways showing the cell cycle arrest in fibrotic matrices (REACTOME database).

#### 3.6.3 Extracellular matrix

The fibrotic microenvironment of C2C12 in the pathological matrices also disturbed the extracellular matrix remodeling. Using the Kegg database, several biological pathways related to extracellular matrix synthesis, its organization and hydrolysis were modulated. Kegg analysis showed that cells/ECM interactions were disturbed. First, some membrane receptors interacting with ECM were downregulated. For instance, Integrin alpha 10 that specifically binds collagen 1 in muscle and hyaluronan receptors (CD 44 and RHAMM) were downregulated (Table 2). Second, major components of lamina membrane such as type IV and VI collagens were downregulated. The synthesis of hyaluronan was also affected as the expression of hyaluronidase synthase 1 and 2 was downregulated. Besides, gene expression of tenascin C, key player in cell migration of satellite cells is inhibited. In addition to a reduced ECM production, matrix remodeling was also inhibited with downregulation of key matrix metalloproteinases (MMP9 and MMP14) involved in matrix remodeling during muscle regeneration (Table 2). Last, several enzymes involved in collagen I maturation and crosslinking were upregulated, as for example, lysyloxidase is a key protein of collagen crosslinking (Table 2).

**Table 2:**
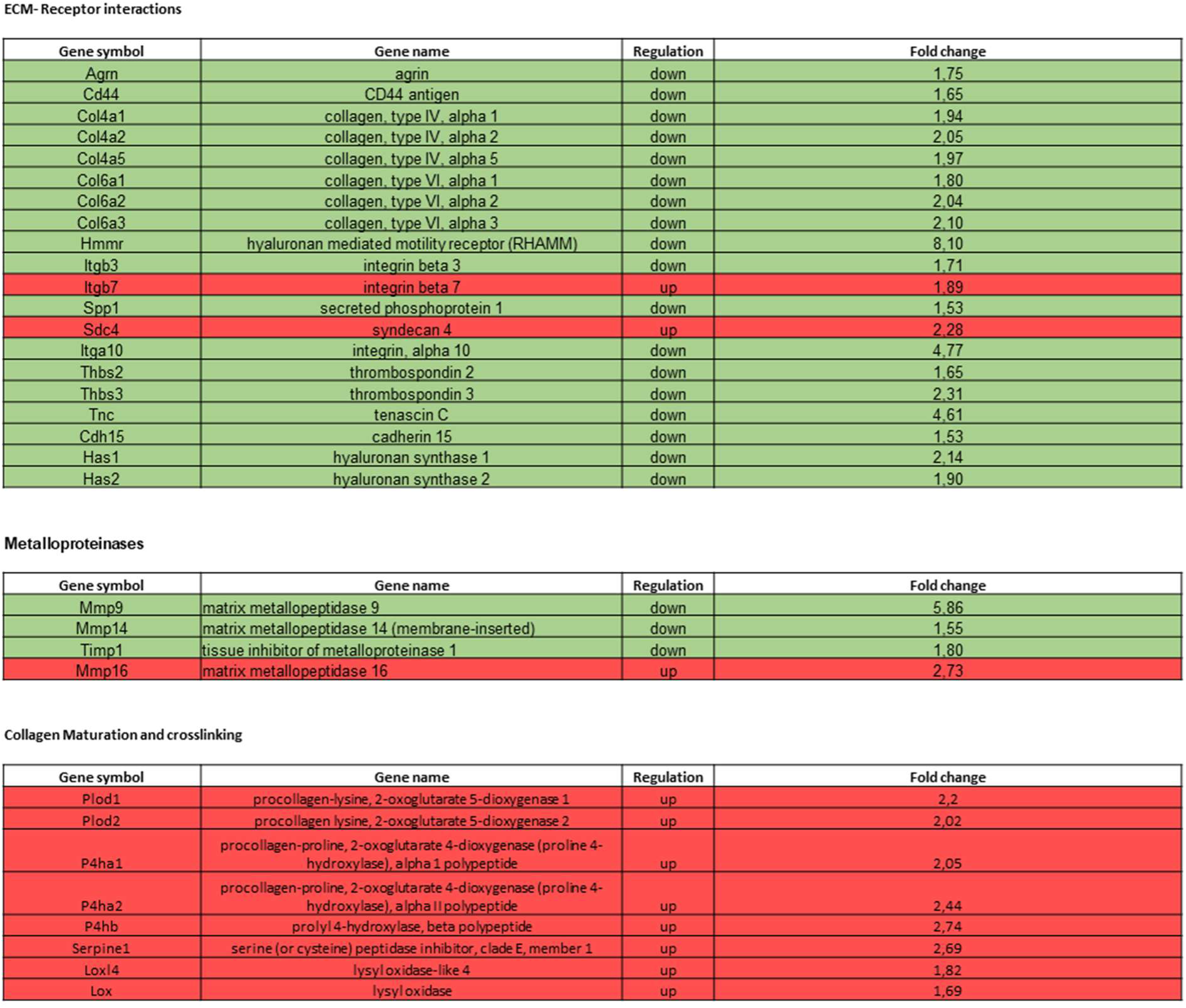
Modulation of genes involved in ECM synthesis, maturation and remodeling. Upregulated genes are in red and downregulated genes are in green.

#### 3.6.4 Myogenesis

Finally, the last crucial aspect of this study consisted in evaluating the myoblast differentiation into myotubes. The RNA sequencing and the GSEA analysis enrichment revealed a diminished myogenesis in pathological matrices (Figure 11 A and B). When the gene expressions were studied in detail, myosin (Myh 1, Myh 3, Mylpf) and troponin (Tnnt1, Tnnt2, Tnnt3, Tnni2) genes were downregulated (Figure 11 C). These proteins are responsible for cell contraction in muscle tissue. Desmin (des) and myoglobin (Mb) were also downregulated, evidencing again an impaired differentiation.

**Figure 11:**
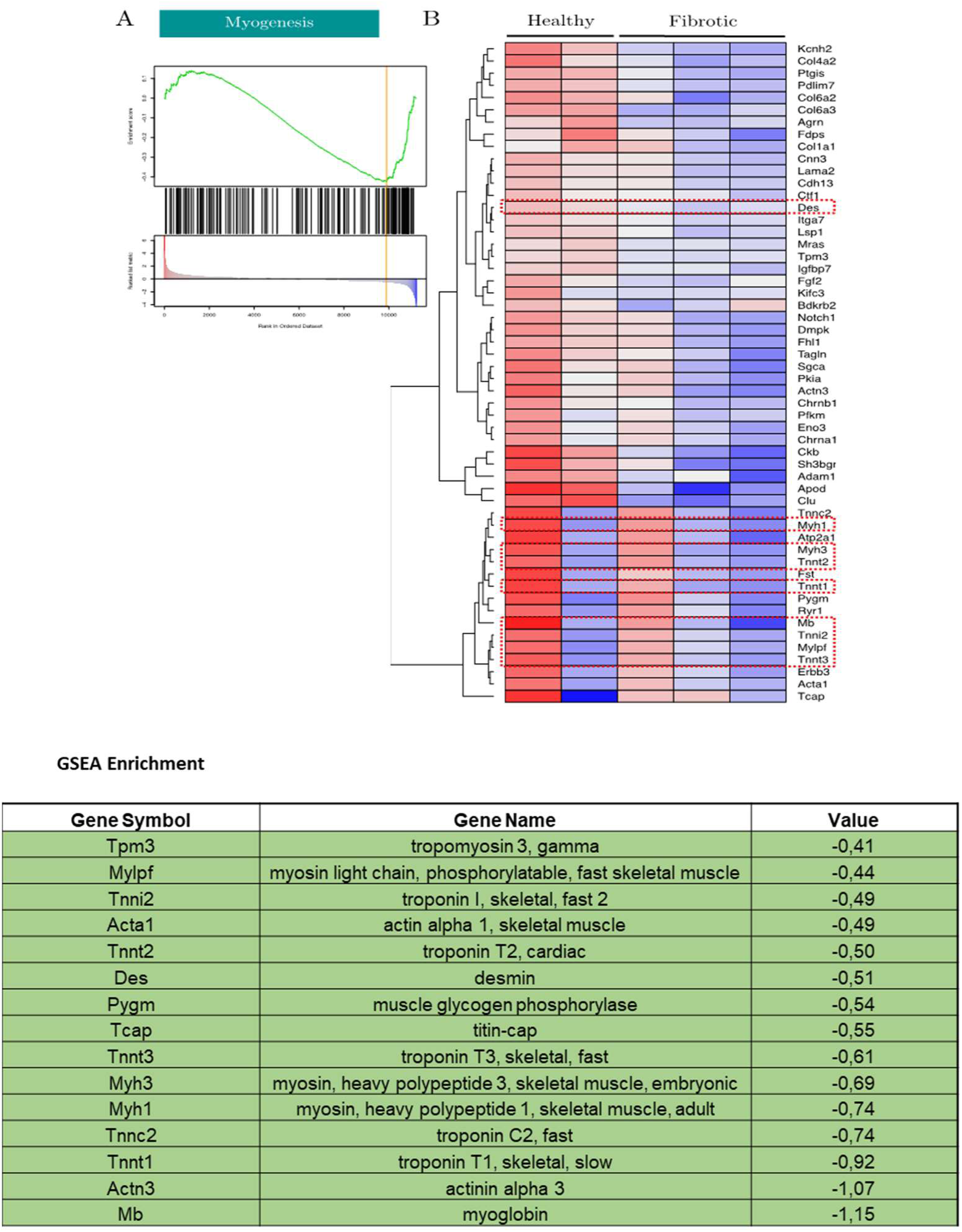
Effect of fibrotic matrix on differentiation of C2C12 myoblasts into mature myotubes. (A): Enrichment analysis of myogenesis gene set, (B) Heat map of regulated genes of the myogenesis gene set in fibrotic and Healthy matrices, (C) list of regulated genes involved in myogenesis. The “Value” column corresponds to the fold change.

### 3.7 Impact of porosity and stiffness on myoblast morphology and differentiation

In addition to the healthy and fibrotic matrices, two supplementary matrices were designed to identify the predominant physical parameter influencing cell behavior within fibrotic matrices. A stiff, porous matrix was created by modifying the healthy matrix through EDC/NHS crosslinking, which increased its stiffness to approximately 50 kPa. Conversely, a soft, non-porous matrix was developed using the same process as for the fibrotic matrices, but without the crosslinking step. Large mature myotubes were observed in healthy matrices (soft and porous) (Figure 12). Cells expressed myosin heavy chain and several nuclei were present within myotubes, thereby evidencing the myoblast differentiation into mature myotubes. Cells aligned within the channels and displayed an organotypic organization with proper cell-to-cell contacts (Figure 12). Increasing the matrix stiffness had a strong impact on cell morphology and differentiation. In stiff, porous matrices, the cells were smaller, and no labelling of myosin heavy chain was detected. Only the actin cytoskeleton was visible (in green, Figure 12). C2C12 cells were not aligned and only contained one nucleus per cell. Therefore, increased the stiffness impaired the myoblast differentiation into mature myotubes. Myoblasts cultured in soft, non-porous matrices exhibited a morphology and organization similar to those observed in stiff, porous matrices. No myosin heavy chain was detected, cells were mononuclear and not aligned. Unlike cells in stiff, porous matrices, actin cytoskeleton was barely visible and no stress fibers were observed. Lastly, cells cultured in fibrotic matrices (stiff and non-porous) were the smallest ones. Some of them were rounded. C2C12 were randomly organized and showed no characteristics of mature myocytes (Figure 12).

**Figure 12:**
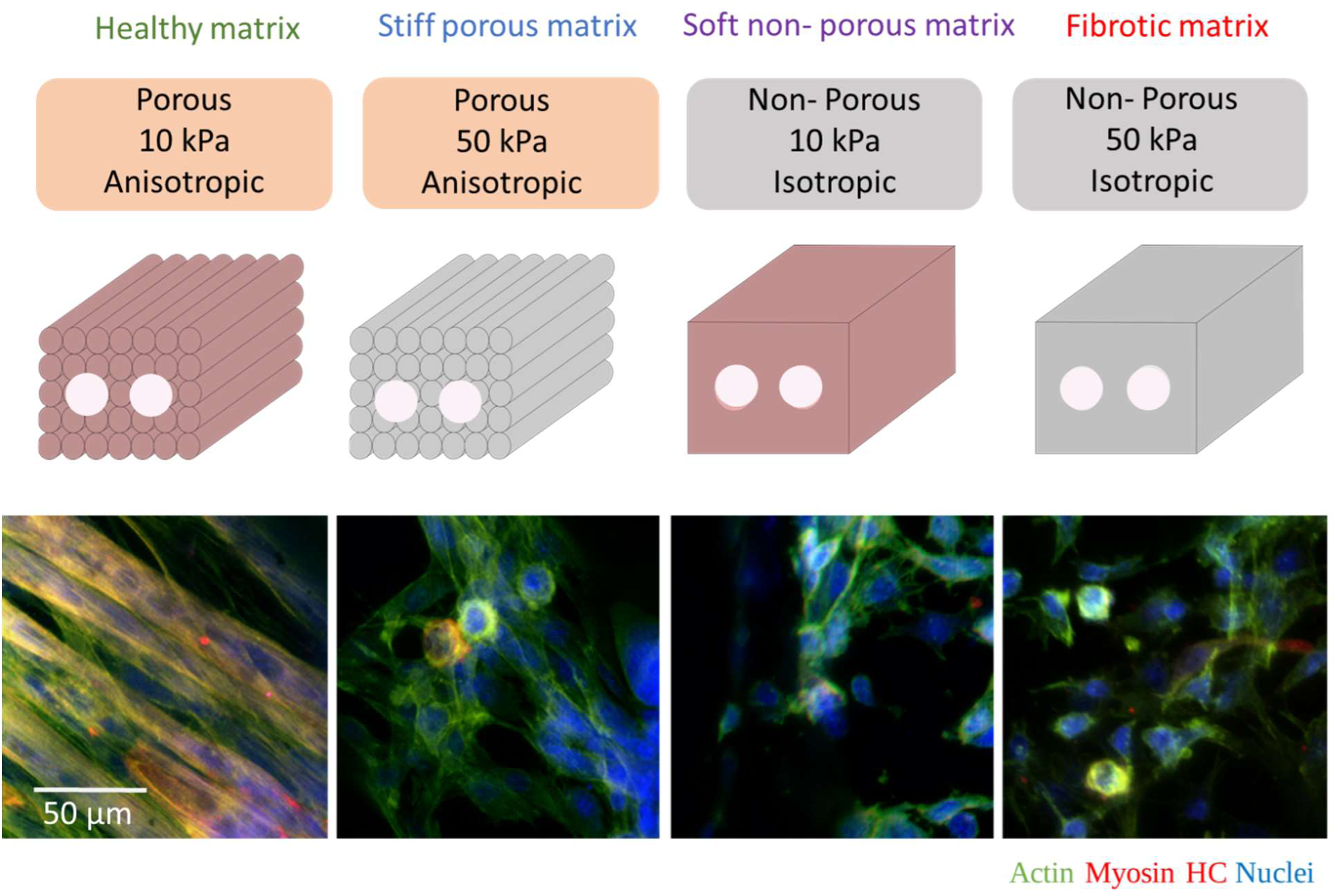
differentiation of myoblasts cultured within healthy matrices, stiff-porous matrices, soft non-porous matrices and fibrotic matrices. Myosin Heavy Chain labeling (red) evidenced cell differentiation into mature myotubes in healthy matrices but not in the other matrices. Actin cytoskeleton was stained in green with Alexa Fluor 488 phalloidin and nuclei were stained in blue with DAPI.

### 3.8 Impact of porosity and stiffness on the cell phenotype of C2C12 myoblasts

#### 3.8.1 Effect of stiffness

Cell culture within stiff and porous matrices triggered the regulation of 790 genes in C2C12 cells, 531 genes were upregulated and 251 were downregulated. The Gene Set Enrichment Analysis (GSEA) revealed that relevant sets of genes were modulated. First, the genes involved in myogenesis were upregulated (Figure 13). The GO analysis confirmed the myoblast differentiation as the genes encoding for dystrophin, myosin, actinin and actin were upregulated. However, the expression of the markers of differentiation was not correlated to the arrest of the cell cycle despite the culture in differentiation medium. The high stiffness of this matrix promoted cell proliferation as shown by GSEA (upregulation of the E2F targets and the G2/Mitose checkpoint) (Figure 13). This was confirmed by the upregulation of several pathways corresponding to different phases/checkpoints of the cell cycle (Reactome analysis). Besides, the expression of inflammatory molecules was downregulated. For example, the signaling pathway of TNF alpha was clearly downregulated. Angiogenesis was also slightly downregulated (Figure 13) and the targets of hypoxia were not modulated, evidencing cells were not in hypoxic conditions. Only the matrix stiffness impacted their phenotype.

**Figure 13:**
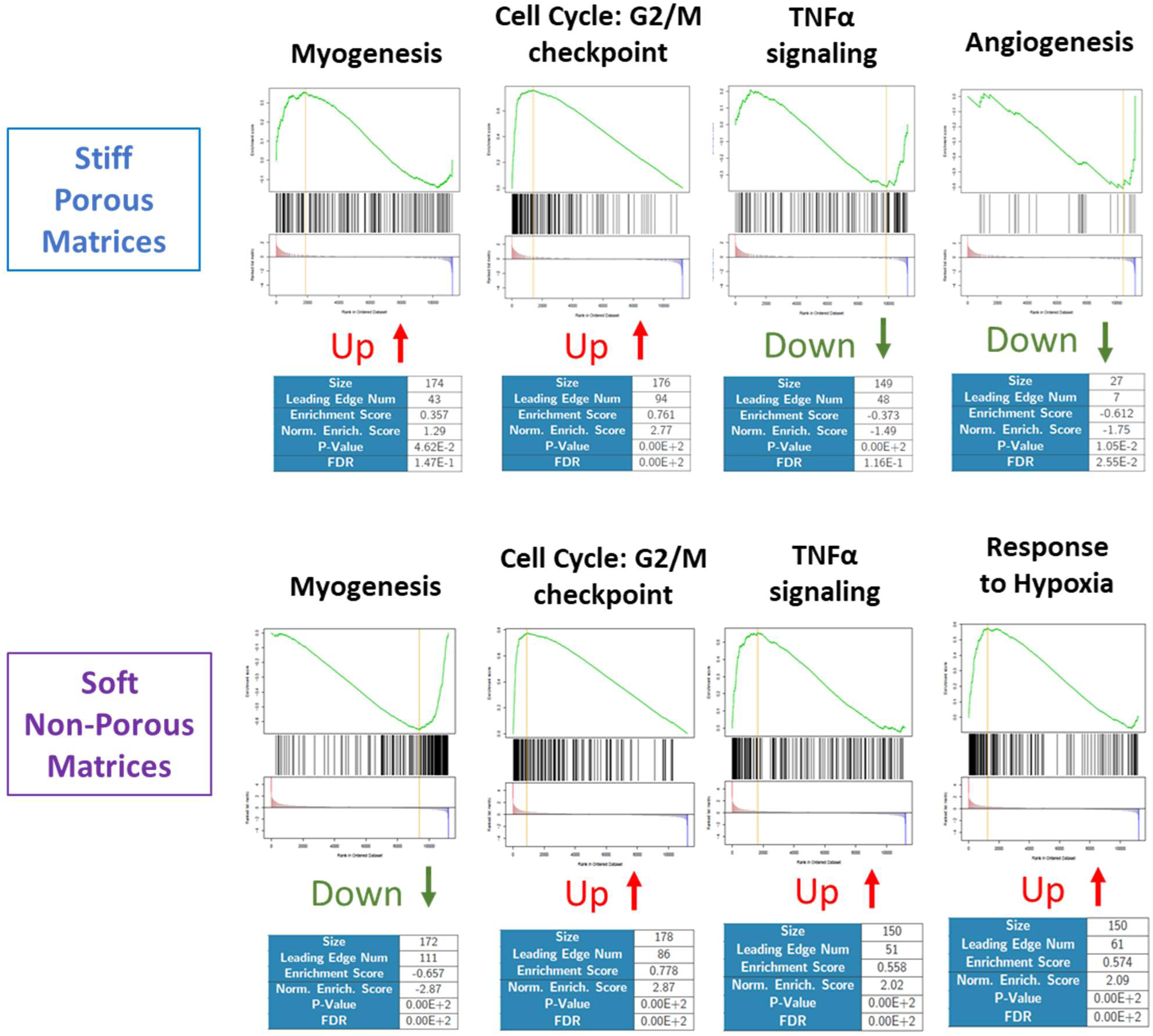
Pathways regulated in stiff porous matrices and soft non-porous matrices analyzed by GSEA.

The bioinformatic analysis revealed that C2C12 myoblasts cultured within stiff matrices upregulated collagen 1 expression and its crosslinking enzymes (LoxL4) (SI N°8). In addition, cells were less prone to remodel their matrix as the major enzymes of degradation (MMP 2, 9, 11 and 13) were downregulated (SI 8). Besides, the cell ability to adhere to collagen was impacted as integrin alpha 10, the major receptor to collagen in skeletal muscle, was downregulated. Last, HA synthase was downregulated whereas laminin, major component of basement membrane, was upregulated (Supplementary Information 9).

#### 3.8.2 Effect of porosity

Myogenesis was significantly inhibited in soft, non-porous matrices, despite being cultured in differentiation medium. GSEA revealed a leading edge number of 111 genes in a set of 171 (Figure 13). Among them, myogenin and myogenic differentiation 1, two transcription factors involved in muscle repair, were downregulated. In contrast, C2C12 proliferation was stimulated in these matrices, as the genes involved in the cell cycle pathway were upregulated (111 genes, Reactome database). These results were confirmed by GSEA (Figure 13). The GSEA showed that genes related to the response to hypoxia were upregulated in soft, non-porous matrices (Figure 13). This evidences the absence of porosity triggers hypoxic conditions in C2C12 cells. Hypoxia was associated with the promotion of angiogenesis as shown by the GO analysis (Supplementary Information 10). Genes involved in inflammation were upregulated in these matrices. For instance, TNF-α signaling pathway was upregulated (GSEA, Figure 13). Unlike stiff, porous matrices, collagen synthesis and crosslinking seems to be downregulated (S.I 9). Overall, the expression of molecules constituting the extracellular matrix or basement membrane were downregulated. Among them, collagen VI, a crucial component of muscle basement membrane, was downregulated (S.I 9). Last, the interactions with ECM and remodeling activities were downregulated (MMP2 and MMP15).

To determine whether stiffness or porosity predominantly influences cell phenotype in fibrotic matrices, Table 3 summarizes key characteristics of C2C12 myoblast phenotype in the different types of matrix. Myogenesis appears to be inhibited under hypoxic conditions due to the lack of porosity. Non-porosity also plays a major role in triggering inflammation and promoting angiogenesis in fibrotic matrices. Surprisingly, the cell cycle was inhibited in fibrotic matrices whereas it was activated in the other ones (stiff, porous and soft non-porous). Stiffness seems to be predominant in the matrix remodeling as the expression of numerous MMPs was downregulated in stiff, porous matrices. In addition, these matrices promote the collagen crosslinking. Lastly, stiffness and non-porosity seems to act in synergy to downregulate the synthesis of basement membrane and promote the switch towards glycolysis.

**Table 3:** Impact of the different matrices on the cell phenotype.

|  | Soft Non-Porous Matrices | Stiff Porous Matrices | Fibrotic Matrices |
| --- | --- | --- | --- |
| Cell Cycle | + | + | - |
| Myogenesis | - | +/- | - |
| Hypoxia | + | - | + |
| Inflammation | + | - | + |
| Glycolysis | + | + | + |
| Angiogenesis | + | - | + |
| Collagen/<br>Crosslinking | - | + | + |
| Basement<br>Membrane<br>synthesis | +/- | +/- | - |
| Remodeling –<br>MMP expression | +/- | - | - |

## 4. Discussion

The primary goal of this study was to design a fibrotic matrix that replicates key structural and physical characteristics of the pathological matrix observed in muscular dystrophies and other degenerative and inflammatory muscular diseases. First, we demonstrated that fibrotic matrices, compared to healthy matrices, lost intrinsic porosity and anisotropy, which are important characteristics of healthy extracellular matrix (ECM). Additionally, fibrotic matrices exhibit significantly higher stiffness [34]. The aim of developing such non-porous and rigid matrices was to recreate the cellular microenvironment typical of dystrophies.

The cell phenotype of myoblasts encapsulated within these pathological matrices was compared to that of cells encapsulated in healthy matrices. Analyzing the impact of the fibrotic matrix on cell behavior is crucial because the muscle cells are not the only actors in the pathophysiology of dystrophies. Besides the impaired cell contraction due to the genetic mutation, the fibrotic matrix gives pathological stimuli form the environment to muscle cells [19].The pathological matrix exhibit specific features such as reduced porosity, high stiffness and disrupted anisotropy [19].

In this study, the 3D printing parameters previously used to synthesize healthy matrices were adapted to mimic the physical properties of pathological muscle matrices [23]. The 3D printing in air associated with a slow gelling by ammonia vapors generated isotropic and non-porous collagen 1 matrices. Collagen molecules extruded during 3D printing aligned thanks to their high concentration, but such collagen concentration is not high enough to maintain molecular alignment like in nematic liquid crystals of collagen [26]. Therefore, collagen molecules relax if the gelling is slow and isotropic dense collagen hydrogels are produced. Nakayama and collaborators have shown in 2D the importance of anisotropy for the muscle cell alignment with the absence of cell organization on isotropic materials [35]. The second consequence of a slow gelling is the spreading of collagen filaments that fulfilled the intrinsic porosity to generate non-porous materials. Hence, by adapting the gelling conditions, matrices with the same collagen concentration exhibited different structural features, *i.e* porous and anisotropic with a rapid gelling and isotropic and non-porous with a slow gelling.

Reproducing the mechanical properties of pathological collagen matrices is a challenging task. Hydrogels fabricated from natural polymers exhibit poor physical properties, far from those observed in skeletal muscle [36]. 3D models are usually made from low concentrated collagen, gelatin or fibrin solutions [15,17,21]. As a consequence, the hydrogel stiffness is very low. To circumvent this drawback, crosslinkers can be used such as UV in GelMA (Gelatin Methacrylate) hydrogels. The stiffness of healthy matrices (12 kPa) can be reached by crosslinking but the latter negatively impacts cell morphology and viability. In addition, these matrices are brittle. In these systems, encapsulated cells do not spread and differentiate [37]. 3D printing of dense collagen concentrated at 30 mg.mL^-1^ enables to reach the stiffness of healthy muscle matrices without crosslinking [23]. Myoblasts are subsequently seeded in the large pores dedicated to cell colonization to allow them to adopt an organotypic organization in fascicles.

Reproducing the fibrotic characteristics of pathological matrices is not feasible using dense collagen solutions concentrated at 30 mg.mL^-1^. The fibrotic matrix is made of highly concentrated collagen 1 (200 mg.mL −1). Such collagen concentration is responsible for the mechanical properties of the pathological matrix. Obtaining such concentrated solution is not technically feasible in vitro due to its high viscosity. In addition, this solution is not printable. As a consequence, mimicking the biochemical characteristics of the fibrotic is not realistic. However, the biochemical nature of such ECM can preserved thanks to the utilization of collagen 1. Nevertheless, it is possible to mimic the physical properties of the fibrotic matrix.

As fibrotic matrices are highly rigid (E=50-70 kPa) and barely deformable, a crosslinking step of 3D printed dense collagen hydrogels could be interesting. The biochemical nature of muscle ECM would be preserved and the physical properties of the pathological tissue reproduced. One strategy based on functionalization of collagen molecules followed by UV crosslinking is not suitable as it affects fibrillogenesis [38]. Glutaraldehyde could be used as it is efficient to reach such mechanical properties but can be toxic for cells. So, the coupling agents EDC/NHC at concentration not toxic for cells were used to reach the physical properties of fibrotic matrices [39]

In vivo, skeletal muscles are organized into fascicles from 400 μm to 5 mm in diameter. These bundles include several myotubes densely-packed together. The final geometry of such organization looks like a cylinder. Large cylindrical pores within healthy and fibrotic matrices had to be created in hydrogels to allow cells to adopt this peculiar morphology. The most popular technique described in the literature consists in introducing needles inside the hydrogel. After gelling and their removal, straight cylindrical pores are obtained [40,41]. Despite its simplicity, this technique reproduces the adequate geometry to allow the muscle fascicle formation after cell seeding and consequently we used this procedure. Cells were mixed with Matrigel® and inject in pores generated by needles in order to reproduce their interaction with their basal lamina. Myoblasts filled the channels after 5 days, remodeled their matrix, organized in 3D and felt the physical properties of their surrounding matrix. An altered differentiation of C2C12 myoblasts was observed in fibrotic matrices. Despite the confinement that should align myotubes [37], the pathological matrix did not allow the generation of muscle fascicles. The reduced differentiation could also be explained by the lack of anisotropy known to induce cell differentiation and proliferation [42]. Current in vitro models developed in the literature never reproduce the pathological conditions. The different systems aim to synthesize matrices possessing the characteristics of healthy muscle ECM. They developed non-porous anisotropic matrices composed of fibrin. Cells located at the hydrogel center suffered from hypoxia and died [20]. Different strategies were tested to increase matrix porosity or vascularization. However, matrices with a high stiffness have not been developed yet and its impact on cell behavior not analyzed. Hence, developing two types of matrices recapitulating the healthy and fibrotic features of muscle ECM seems to be of interest to study the impact of the pathological matrix on cell phenotype.

Hypoxia is a main feature of muscular dystrophies [43]. To evidence that cells were in hypoxia in fibrotic matrices, several markers have been identified. It is well established that the accumulation of HIF-alpha, a key molecule in hypoxia, in C2C12 cells leads to the upregulation of angiogenic factors. In fibrotic matrices, the expression of VEGF-A, -B, and -D is increased, suggesting reduced oxygen levels in the vicinity of the cells [44]. HIF-alpha also triggers the shift from mitochondrial respiration to pure glycolysis and production of reactive oxygen species [45]. The gene expression study revealed several enzymes involved in glycolysis and ROS production were upregulated, thereby evidencing cells are in hypoxia. In tissues, cells located more than 200 µm from capillaries suffer from hypoxia. In fibrotic matrices, C2C12 cells have been seeded in central channels with a low nutrient and O2 diffusion. The culture medium needs to diffuse approximately 700 µm to reach the cells. In addition, high collagen concentration is correlated with slow liquid and O_2_ diffusion [24,46]. Last, fibrotic matrices are cross-linked to increase their stiffness. This could decrease the gas diffusion through matrices.

The role of hypoxia on muscle cell proliferation is still under debate. Some groups showed that hypoxic conditions accelerate regeneration after muscle injury (proliferation and differentiation) but others observed that a low O_2_ level induces quiescence of satellite cells [47,48]. Interestingly, inhibition of proliferation could depend on the degree of hypoxia. Beaudry and collaborators showed that cells proliferate when cultivated under 5% O_2_ but are quiescent when the O_2_ level decreased down to 1% [49]. In fibrotic matrices, the downregulation of numerous genes involved in the cell cycle suggests that the absence of porosity would have a negative impact on cell proliferation. In addition, hypoxia seems to be severe (less than 1%). Interestingly, C2C12 cultured in non-porous but not cross-linked matrices suffers from hypoxia as revealed by RNA sequencing but their cell cycle is not arrested. We can hypothesize that the degree of hypoxia is lower and allow cell proliferation. Hence, the matrix stiffness could increase hypoxia in C2C12 cells by decreasing O_2_ diffusion.

In pathological matrices, C2C12 cells do not differentiate into mature myotubes. Moreover, they do not align and do not adopt the physiological 3D organization. Hence, the cell cycle arrest was not correlated with cell differentiation into myotubes. Hypoxic conditions could be at the origin of an impaired differentiation towards the adipogenic lineage as suggested by the GSEA analysis [50,51]. Last, hypoxic conditions could also explain the expression of inflammatory cytokines, which is a characteristic of muscular dystrophies [52,53]. This is supported by the cell culture in soft non-porous matrices where C2C12 are in hypoxia and expressed inflammatory cytokines.

Fibrotic collagen matrices are also characterized by their high stiffness. The latter modulates cell adhesion on their matrix and regulates myogenesis [54,55]. Some groups have shown that the physiological rigidity around 12 kPa is required to promote adequate differentiation but stiffer or softer substrates do not promote appropriate myogenesis [55,56]. In our condition, the high stiffness promotes the gene expression of myogenesis markers. However, cells cultured in stiff and porous matrices did not differentiate into mature myotubes as seen by confocal microscopy. In addition, they did not organize in fascicles. Hence, hypoxia and stiffness seem to act in synergy to inhibit the myoblast differentiation into myotubes. Actually, cell morphology seems to be more altered in fibrotic matrices compared to matrices with a single characteristic as they are rounder and smaller.

Matrix stiffness also regulates the adhesion strength and the cell/ECM interactions [57]. In fibrotic matrices, many genes involved in collagen stability and its natural cross-linking are upregulated. As the matrix is already very stiff, this can generate a vicious cycle as cells promote matrix stiffening. Using stiff, porous matrices, we confirm that stiffness is the main stimulus to rigidify matrices by increasing collagen crosslinking and production.

The biochemical composition of the basal lamina is the basis of regeneration of damaged muscle as abnormalities in its structure or biochemical composition is associated with impaired myofiber integrity and muscular dystrophy [19,38]. In fibrotic matrices, components of lamina such as collagen IV and VI were downregulated. Therefore, contacts between myoblasts and lamina are altered and do not allow cell fusion [40,58,59]. Hence, the biochemical and biophysical properties of the fibrotic matrix impaired its remodeling by cells. This is confirmed by the downregulation of MMP14 and MMP9, two majors enzymes involved in matrix hydrolysis, which allow cell migration to the injury site and matrix remodeling [60,61]. Therefore, cells in fibrotic matrices are unable to restore a physiological basal lamina required for muscle regeneration. Using the matrices possessing a single characteristic of the pathological ECM, it was shown that stiffness was predominant to inhibit degradation whereas the absence of porosity had a detrimental effect on the expression of the basement membrane proteins. However, both parameters were detrimental for the matrix remodeling.

The altered cell/matrix interactions also impact hyluronan (HA) metabolism. The enzymes involved in HA synthesis and degradation were downregulated. In addition, HA receptors were also downregulated in fibrotic matrices. As hyaluronan and its receptors plays an important part in myogenesis, this could partially explain the altered differentiation observed in fibrotic matrices [62,63].

## 5. Conclusion

In this study, healthy and fibrotic matrices were synthesized to analyze the impact of the pathological microenvironment on cellular behavior in muscular dystrophy. Through 3D printing, adequate gelling conditions and crosslinking step, non-porous, isotropic and rigid hydrogels were generated mimicking the features of pathological muscle ECM. C2C12 murine myoblasts seeded within matrices sensed their microenvironment, exhibiting distinct behaviors based on the matrix’s physical properties. The fibrotic matrix inhibited cell proliferation and differentiation into mature and elongated myotubes. In addition, 3D cell organization was also altered. Gene Set Enrichment Analysis (GSEA) revealed that the cells were in hypoxic conditions, had a diminished ability to remodel the ECM, and secreted inflammatory cytokines. These findings suggest that the fibrotic matrix in dystrophies hampers myoblasts’ ability to regenerate muscle. Therefore, developing two relevants 3D models—one healthy and one fibrotic— provides a valuable platform for studying the pathophysiology of muscular dystrophies, specifically the influence of the persistent fibrosis on progression of muscle dysfunction.

## Acknowledgements

The authors thank PhD. Lotfi Slimani for micro-computed tomography imaging from the Life Imaging Facility of Paris University supported by France Live Imaging (grant ANR 11-INBS-0006) and Infrastructures Santé. We also thank Korin Ozkaya and Blaise Hébert for their help.

## Sample CRediT author statement

**Marie Camman**: Conceptualization, Methodology, Formal analysis, Investigation, Writing - Original Draft. **Naomi Nieswic**: Investigation, Methodology. **Pierre Joanne**: Methodology, Conceptualization, Writing - Review & Editing. **Julie Brun**: Methodology, Investigation. **Alba Marcellan**: Conceptualization, Validation, Writing - Review & Editing. **Onnik Agbulut**: Funding acquisition, Supervision, Validation, Writing - Review & Editing. **Christophe Helary**: Funding acquisition, Supervision, Conceptualization, Validation, Methodology, Writing - Original Draft.

## Competiting interests

The authors declare no conflicts of interest.

## Author Approvals

All authors have seen and approved the manuscript. Iit hasn’t been accepted or published elsewhere.

## Supplementary Information

**Supplementary Information 1:**
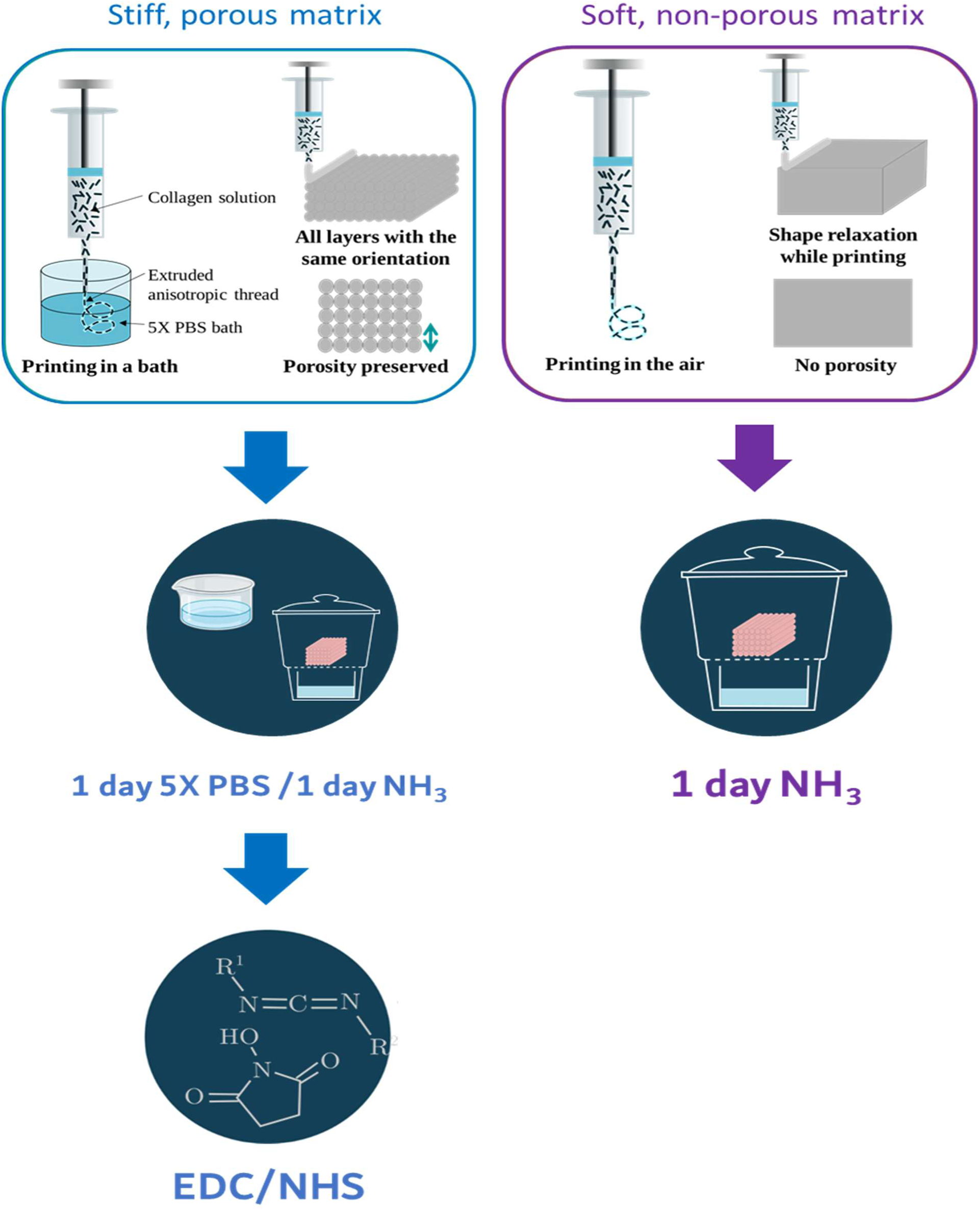
Fabrication of stiff, porous and soft, non-porous matrices by 3D printing. Stiff, porous matrices were printed in 5X PBS, gelled in 5X PBS for one day and in ammonia vapors for another day. After several rinses to decrease the pH down to 7, matrices were crosslinked using EDC/NHS. Soft, non-porous matrices were printed in air and gelled in ammonia vapors for one day.

**Supporting Information 2:**
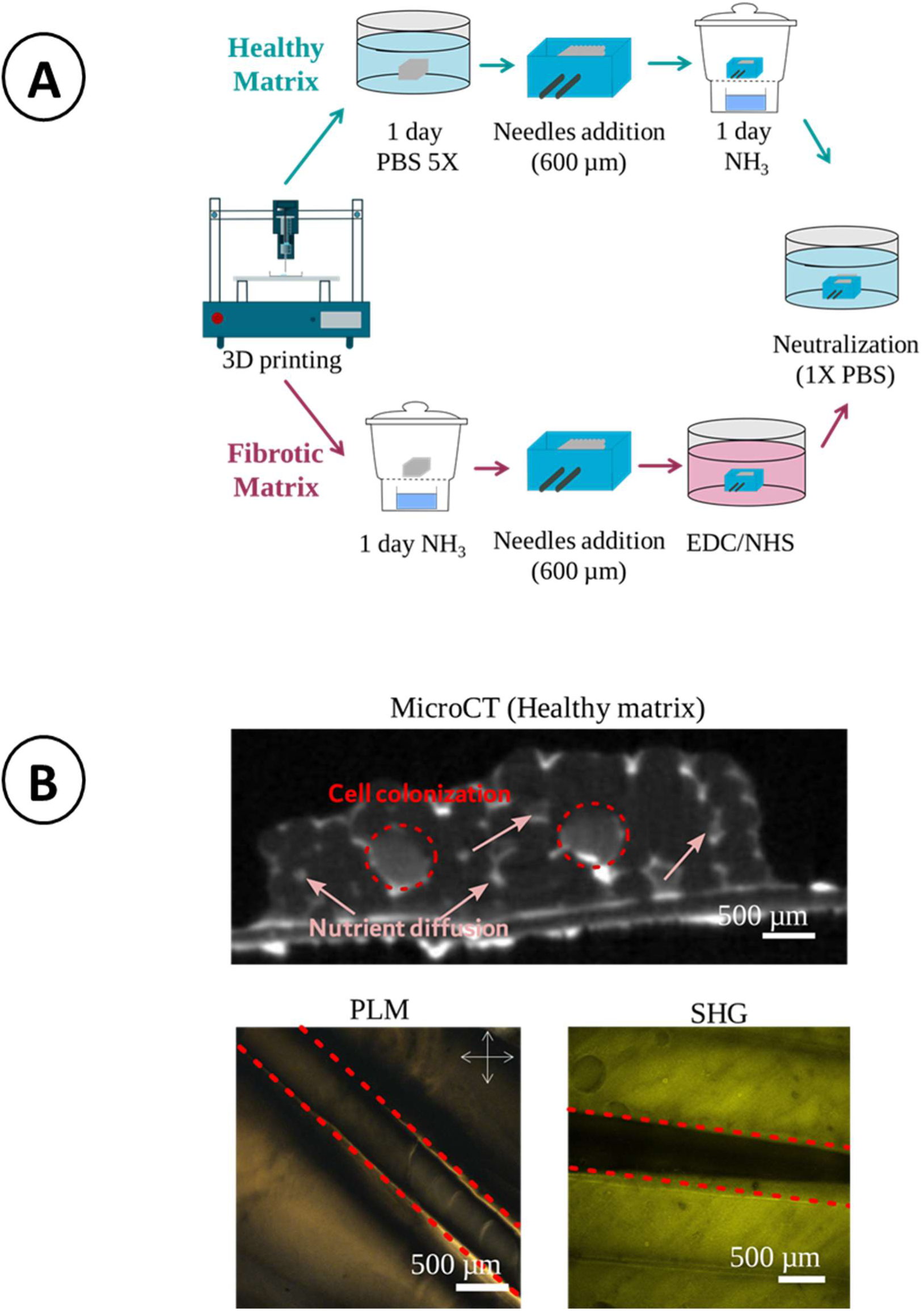
(A) Generation of macroporosity dedicated to cell colonization by introduction of needles (B) macroposoity observed by X X-Rays microtomography (MicroCT), Polarized Light Microscopy (PLM) and second harmonic generation microscopy (SHG).

**Supplementary Information 3:**
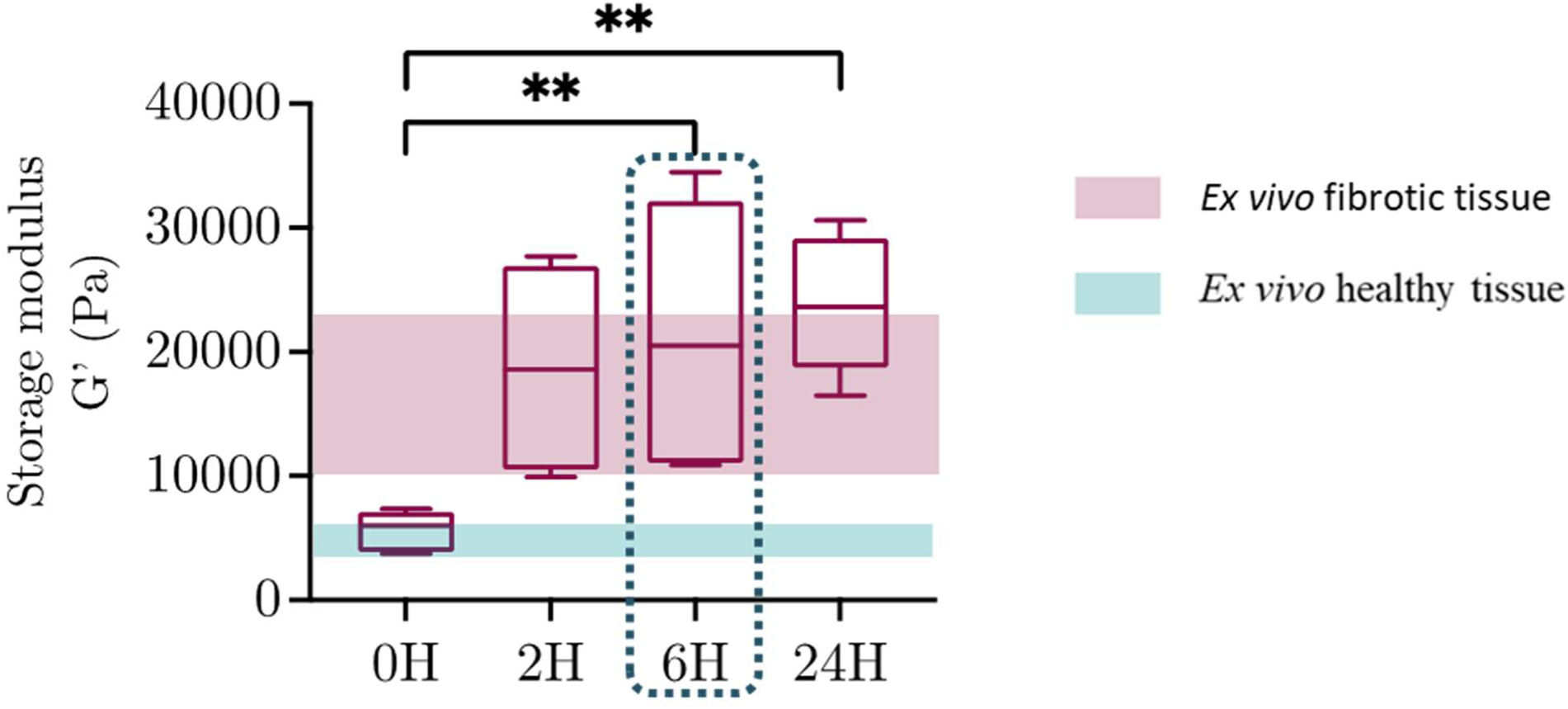
Storage modulus of collagen matrices crosslinked with 5/1.2 mg.mL^-1^ EDC/NHS versus time of crosslinking.

**Supplementary Information 4:**
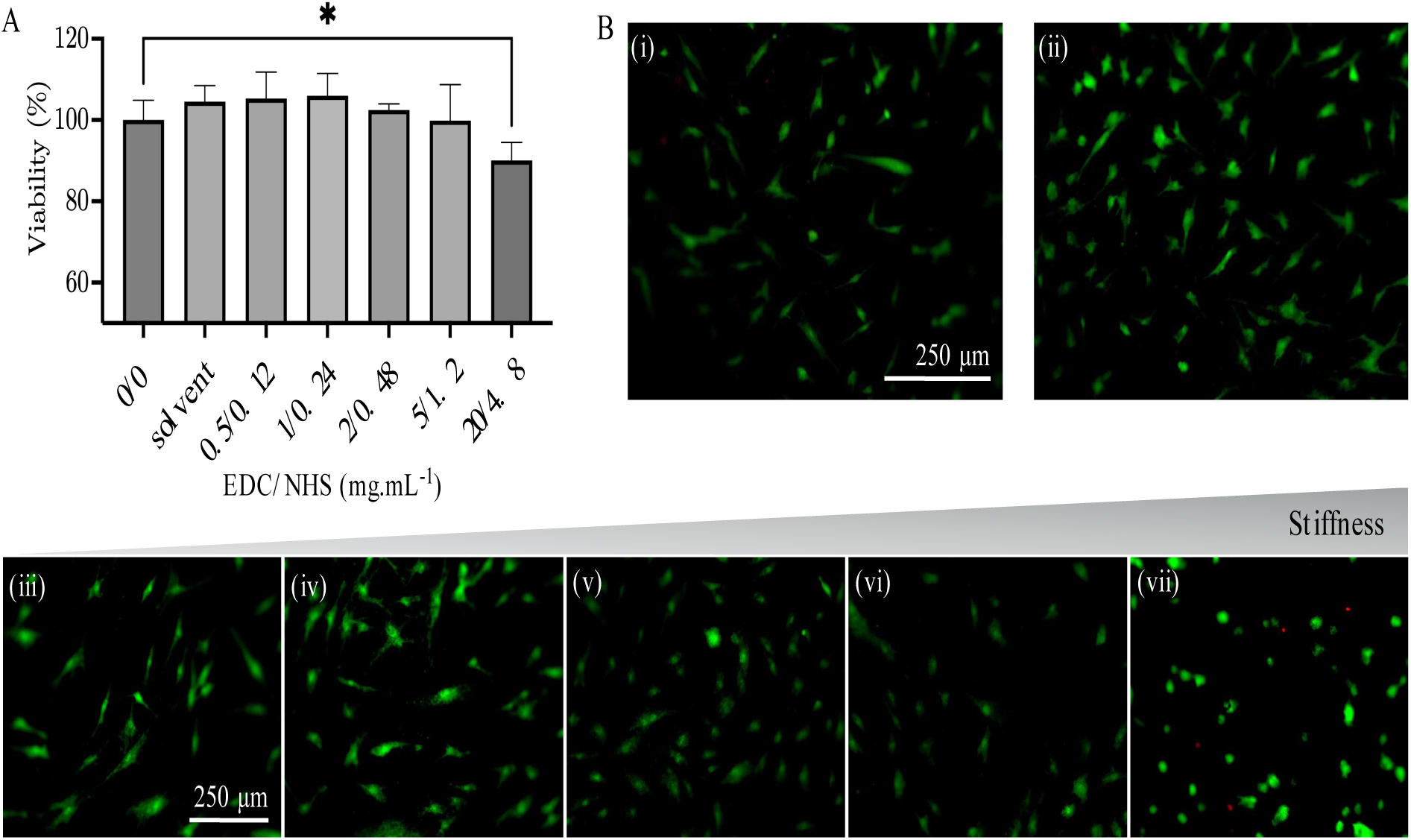
EDC/NHS cytotoxicity on normal human dermal fibroblasts. (A): Alamar blue assay (4 hours) to assess cell metabolism. (B): Live (green)/Dead (red) staining for the different concentrations of crosslinkers. Collagen hydrogel (i) without crosslinker, (ii) with Ethanol 75%, (iii) EDC/NHS=0.5/0.12 mg.mL-1, (iv) EDC/NHS=1/0.24 mg.mL-1, (v) EDC/NHS=2/0.48 mg.mL-1, (vi) EDC/NHS=5/1.2 mg.mL-1, (vii) EDC/NHS=20/4.8 mg.mL-1. *: p<0.05.

**Supplementary Information 5:**
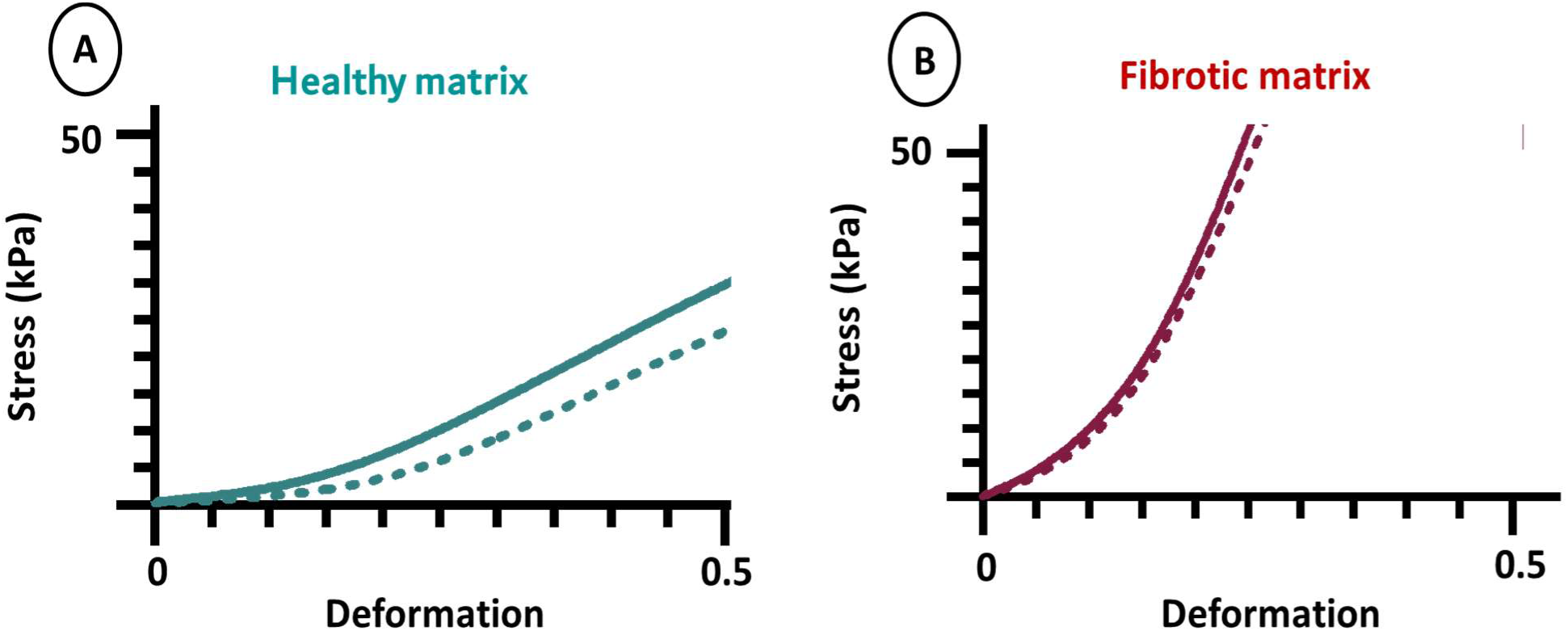
Mechanical properties of 3D printed healthy (A) and pathological matrices (B) assessed by tensile tests. Matrices were longitudinally stretched (plain lines) or perpendicularly stretched (dotted lines)

**Supplementary Information 6:**
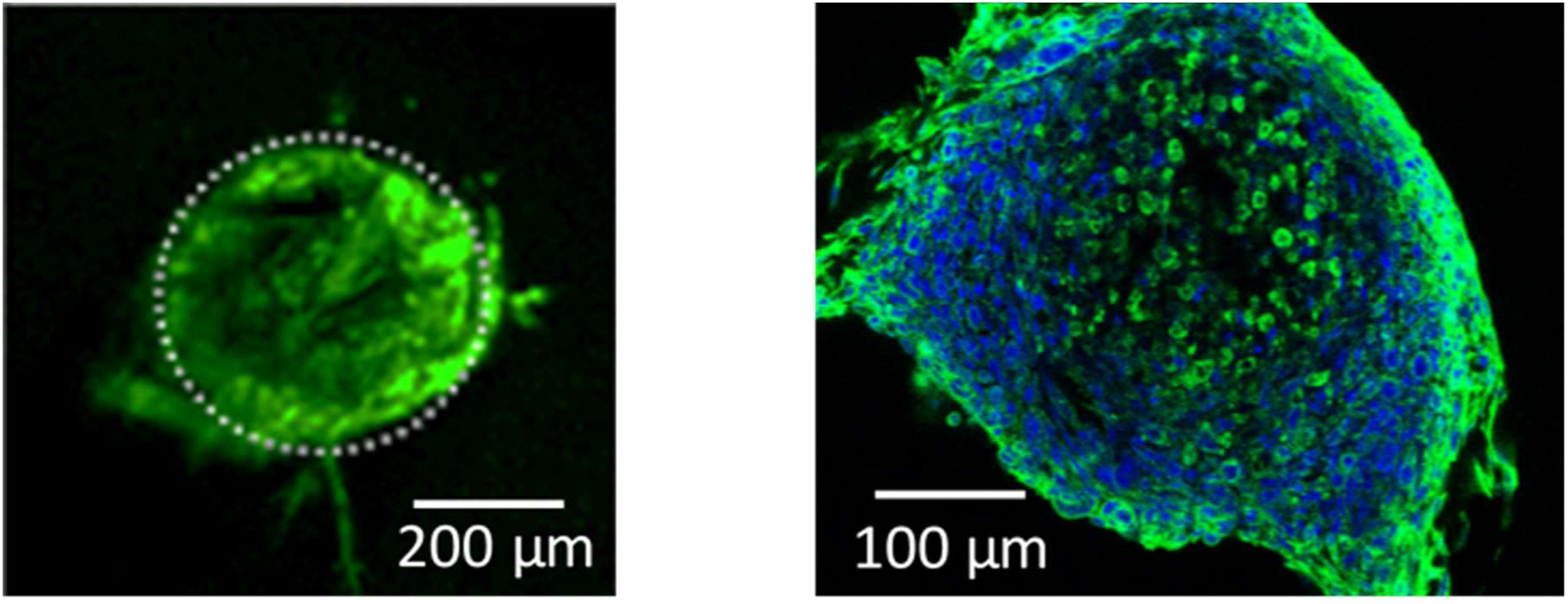
Transversal view of macropores dedicated to cell colonization after 5 days in culture. Nuclei labelledin blue (DAPI) and cytoskeleton in green (Alexa Fluor 488 Phalloidin)

**Supporting Information 7:**
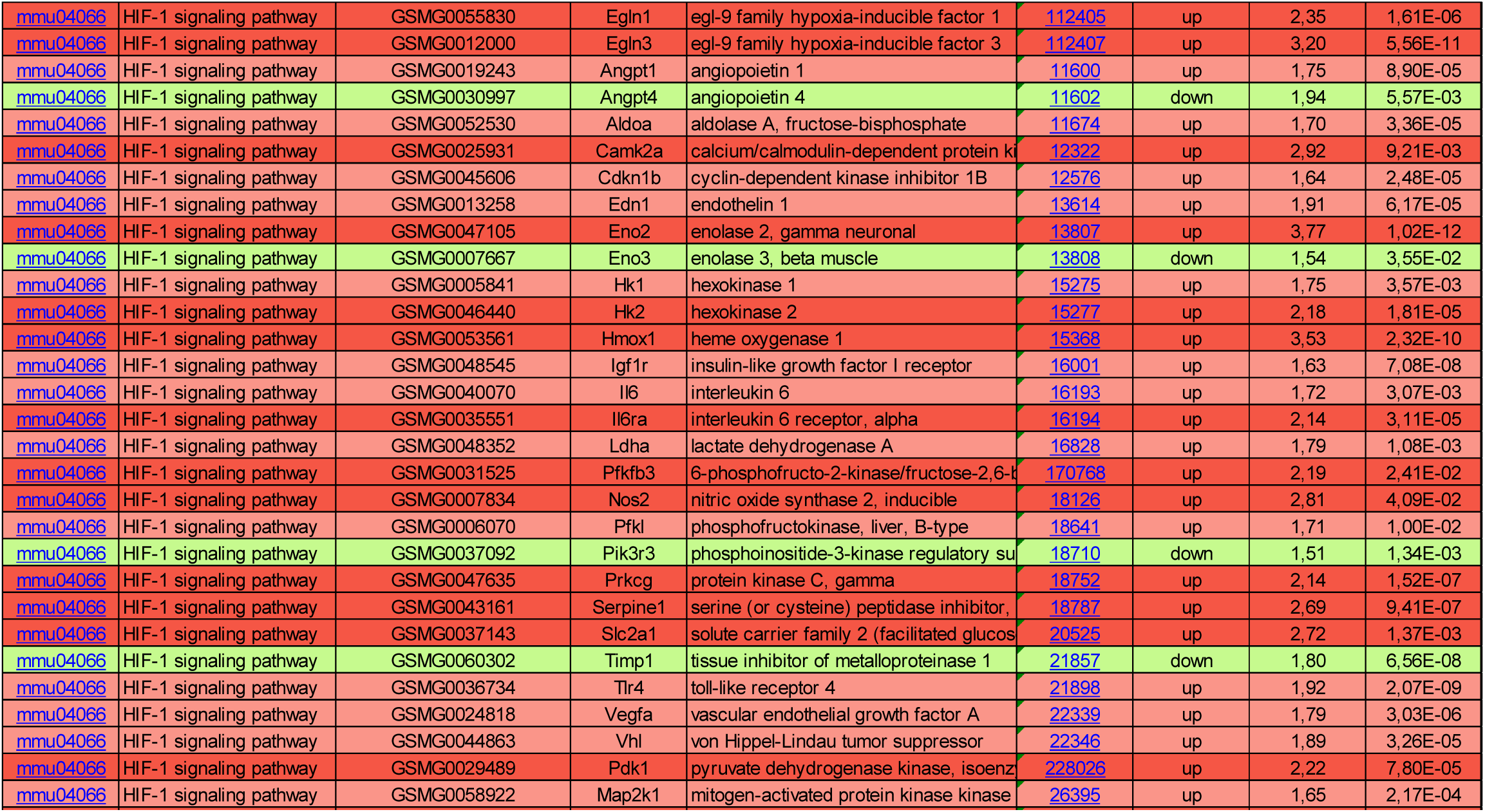
Comparsion Fibrotic matrices vs Healthy matrices. List of upregulated (in red or pink) or downregulated (in green) genes in response to hypoxia.

**Supporting Information 8:**
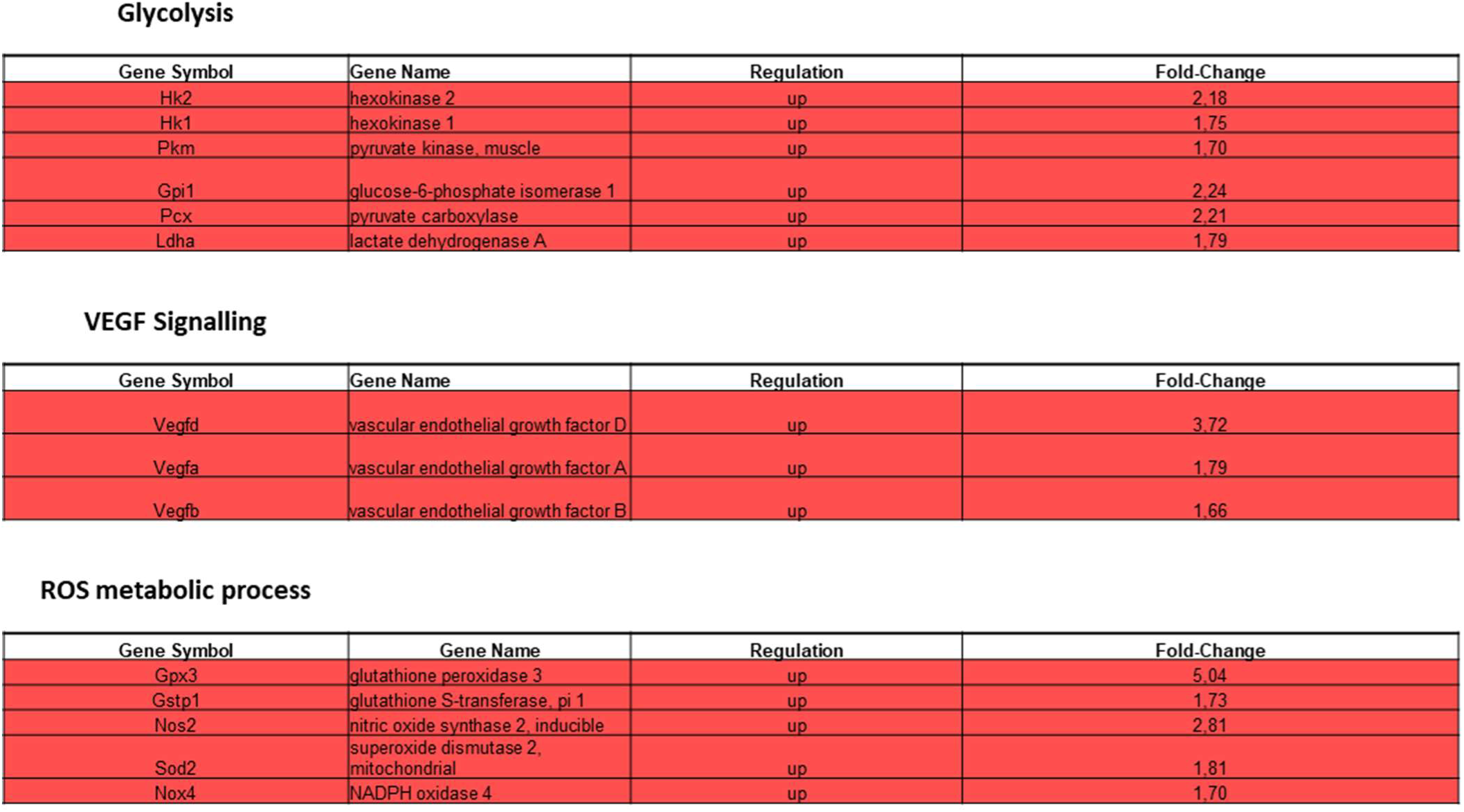
Comparsion fibrotic matrices vs healthy matrices. Regulation of genes involved in glycolysis, VEGF signaling and ROS metabolic process. Upregualted genes are in red.

**Supporting Information 9:**
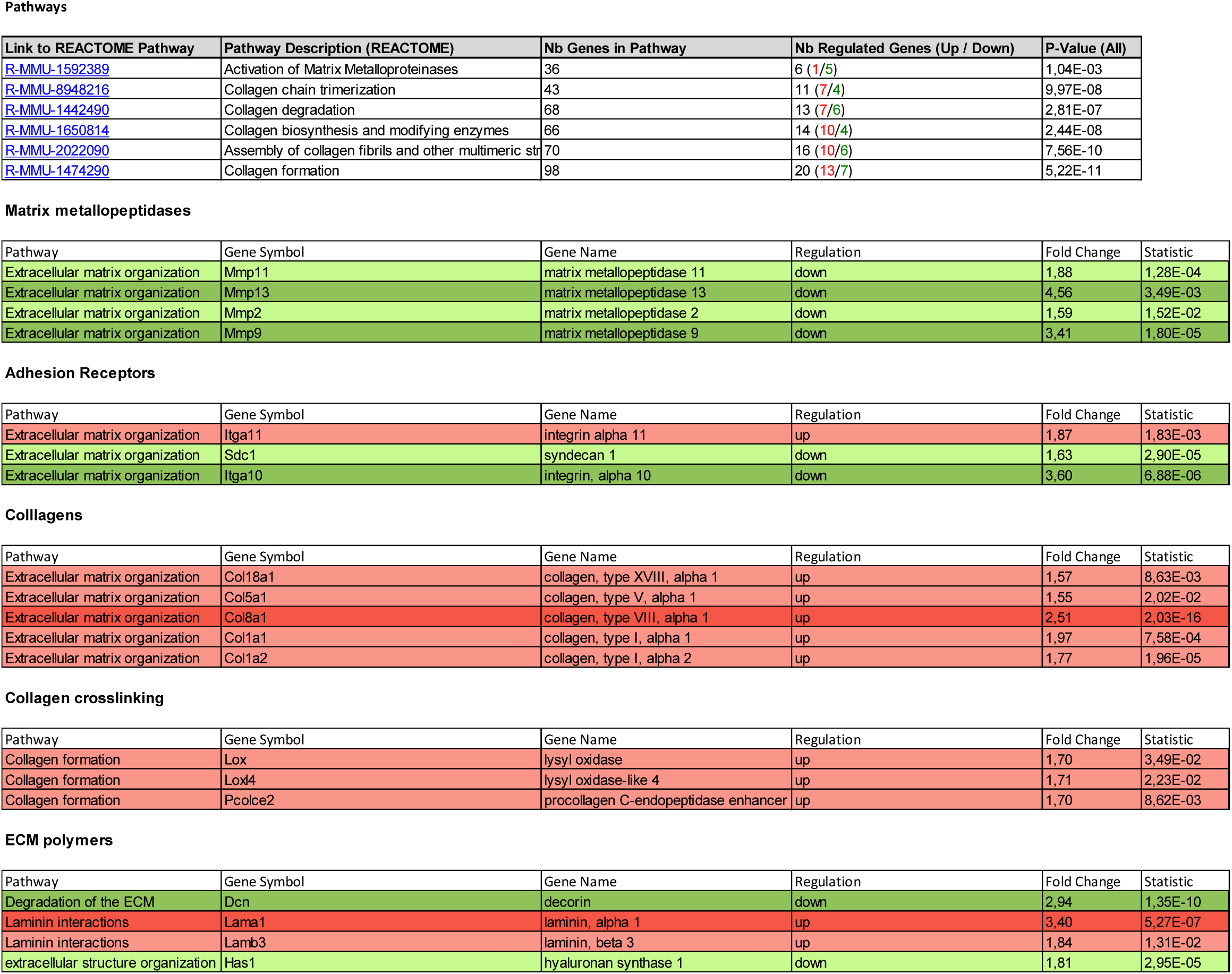
Comparison stiff porous matrices vs healthy matrices. List of relevant pathways regulated (Reactome Analysis). List of upregulated (in red or pink) or downregulated (in green) genes.

**Supporting Information 10:**
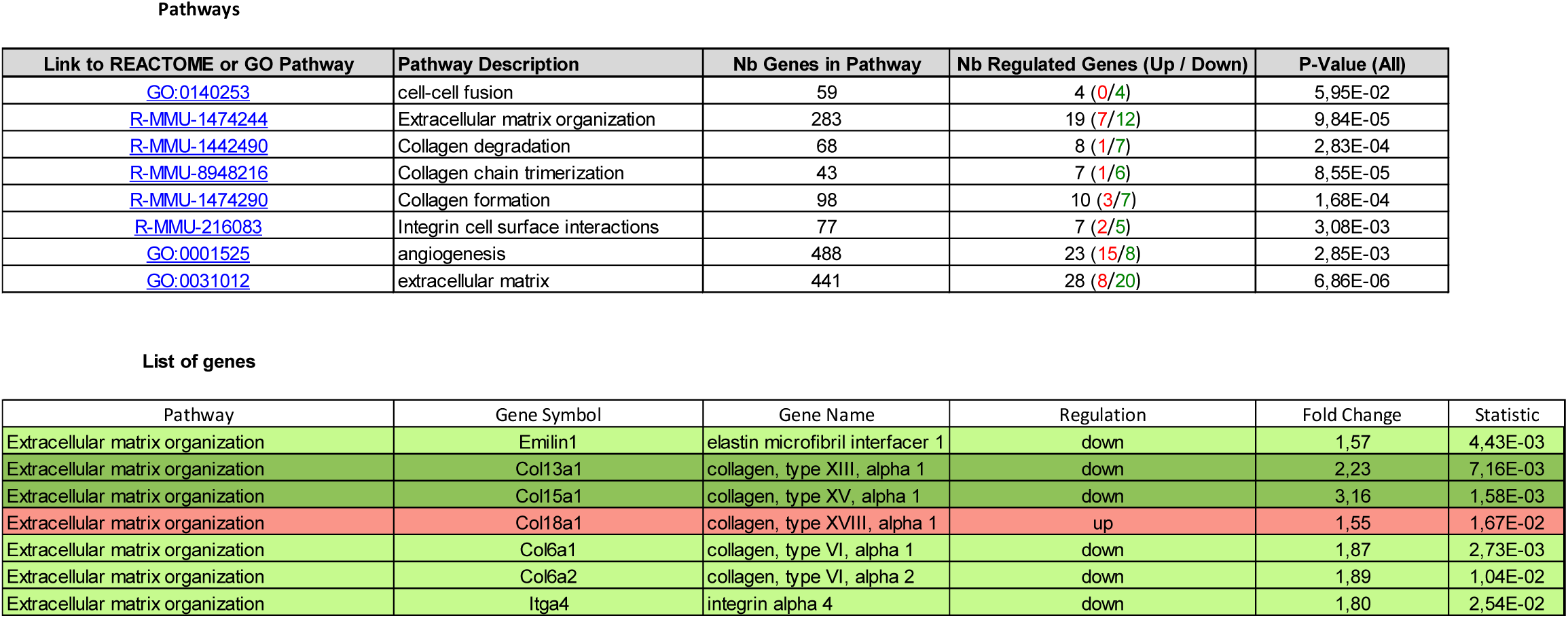
Comparsion soft non-porous matrices vs healthy matrices. List of relevant pathways regulated (Reactome or GO Analysis). List of upregulated (in red or pink) or downregulated (in green) genes.

---

**Table 1:** Cross-linker concentrations tested to rigidify dense collagen matrices.

| EDC | NHS |
| --- | --- |
| 0 mg·mL<sup>-1</sup> | 0 mg·mL<sup>-1</sup> |
| 0.5 mg·mL<sup>-1</sup> (3.2 mM) | 0.12 mg·mL<sup>-1</sup> (1 mM) |
| 1 mg·mL<sup>-1</sup> (6.4 mM) | 0.24 mg·mL<sup>-1</sup> (1.5 mM) |
| 2 mg·mL<sup>-1</sup> (12.8 mM) | 0.48 mg·mL<sup>-1</sup> (3.1 mM) |
| 5 mg·mL<sup>-1</sup> (32.2 mM) | 1.2 mg·mL<sup>-1</sup> (7.7 mM) |
| 20 mg·mL<sup>-1</sup> (129 mM) | 4.8 mg·mL<sup>-1</sup> (30.1 mM) |

